# Single-cell sequencing unveils distinct immune microenvironment with CCR6-CCL20 crosstalk in human chronic pancreatitis

**DOI:** 10.1101/2021.04.05.438347

**Authors:** Bomi Lee, Hong Namkoong, Yan Yang, Huang Huang, David Heller, Gregory L. Szot, Mark M. Davis, Stephen J. Pandol, Melena D. Bellin, Aida Habtezion

## Abstract

**Objective:** Chronic pancreatitis (CP) is a potentially fatal disease of the exocrine pancreas, with no specific or effective approved therapies. Due to difficulty in accessing pancreas tissues, little is known about local immune responses or pathogenesis in human CP. We sought to characterize pancreas immune responses using tissues derived from patients with different etiologies of CP and non-CP organ donors in order to identify key signaling molecules associated with human CP.

**Design:** We performed single-cell level cellular indexing of transcriptomes and epitopes by sequencing (CITE-Seq) and T cell receptor sequencing of pancreatic immune cells isolated from organ donors, hereditary, and idiopathic CP patients who underwent total pancreatectomy. We validated gene expression data by performing flow cytometry and functional assay in the second CP patient cohort.

**Results:** Deep single-cell sequencing revealed distinct immune characteristics and significantly enriched CCR6^+^ CD4^+^ T cells in hereditary compared with idiopathic CP. In hereditary CP, a reduction in T cell clonality was observed due to the increased CD4^+^ T (Th) cells that replaced tissue-resident CD8^+^ T cells. Shared TCR clonotype analysis among T cell lineages also unveiled unique interactions between CCR6^+^ Th and Th1 subsets, and TCR clustering analysis showed unique common antigen binding motifs in hereditary CP. In addition, we observed a significant upregulation of the CCR6 ligand (*CCL20*) among monocytes in hereditary CP as compared with those in idiopathic CP. The functional significance of CCR6 expression in CD4^+^ T cells was confirmed by flow cytometry and chemotaxis assay.

**Conclusion:** Single-cell sequencing with pancreatic immune cells in human CP highlights pancreas-specific immune crosstalk through the CCR6-CCL20 axis that might be leveraged as a potential future target in human hereditary CP.

**Significance of this study:** *What is already known about this subject?:*

- Chronic pancreatitis (CP) is considered an irreversible fibroinflammatory pancreatic disease and remains a major source of morbidity among gastrointestinal diseases with no active approved therapy.
- Inflammation is a known hallmark and contributor to CP pathogenesis. However, little is known about local immune responses in human CP especially with different etiologies.

*What are the new findings?:*

- Single-cell RNA sequencing of pancreatic immune cells from CP patients and organ donors revealed distinct immune transcriptomic features in CP versus non-diseased controls and hereditary versus idiopathic CP.
- Single-cell T cell receptor sequencing unveiled pancreas-specific clonal expansion in CD8^+^ T cells and CD4^+^ T cells-driven unique TCR repertoire changes in hereditary CP.
- We identified that the CCR6-CCL20 axis was significantly upregulated in hereditary CP compared with controls or idiopathic CP suggesting a potential future target for human hereditary CP.

*How might it impact on clinical practice in the foreseeable future?:*

- The results of this study will improve our understanding of CP heterogeneity and identify distinct immune responses in different types of human CP that could provide novel conceptual directions for therapeutic strategies in treating hereditary and idiopathic CP.
- The CCR6-CCL20 axis found in hereditary CP could be a potential novel targetable signaling pathways in the treatment of hereditary CP.

## INTRODUCTION

CP is a progressive fibro-inflammatory disease of the exocrine pancreas and remains a major source of morbidity yet remains an untreatable disease so far^1 2^. CP is characterized by morphological changes in the pancreas including acinar cell atrophy, distorted pancreatic ducts, inflammation, and fibrosis^3 4^. Primary risk factors in adult CP include alcohol and smoking, but genetic variants and idiopathic factors are significant contributors for CP of all ages^1 5-8^. Over the past two decades, many animal models have been used to understand disease pathogenic mechanisms in CP^9 10^. However, questions remain regarding the translational accuracy of pre-clinical studies. For instance, most pancreatitis animal studies are dependent on disease inducing-agents that may not mimic the human CP condition^11^. Therefore, research with human pancreas tissues is critical to understand human disease-specific pathogenic mechanisms in CP.

Inflammation is a hallmark of CP^4 12^, and immune cells have emerged as key contributors in CP and its progression^13 14^. For example, anti-inflammatory (M2) macrophages were observed in different types of CP experimental models^15 16^ and contribute to CP fibrogenesis through the crosstalk with pancreatic stellate cells (PSCs)^17^. A growing number of studies demonstrated the critical role of immune cells at the different stages and disease progression in CP providing potential therapeutic targets for the disease^18-20^. However, immune characteristics remain poorly understood in human CP due to the limited access to human pancreatic tissues, and the contribution of disease etiologies to disease heterogeneity remains unexplored.

In collaboration with an institution that performs high volume of total pancreatectomy islet auto-transplantation (TPIAT) in CP patients, our pilot study using flow cytometric analysis and bulk T cell receptor (TCR) sequencing revealed distinct immune responses between hereditary and idiopathic CP implicating different immunopathogenic mechanisms underlying two different subtypes of CP^21^. Here we performed CITE-seq (combined single-cell antibody-derived tag and RNA sequencing)^22^ and single-cell TCR sequencing (scTCR-seq) of immune cells isolated from human pancreas tissues in CP patients and organ donors for a more in-depth understanding of immune responses associated with the disease. This unbiased systemic analysis of immune signatures that included protein expressions, transcriptomes, and TCR repertoire analyses of pancreatic immune cells revealed distinct and unique disease-specific immune cell signatures and interactions that provide insights into novel immune signaling pathways and offer potential future therapeutic target(s) for human CP.

## RESULTS

### Human pancreatic immune cell transcriptional atlas reveals unique disease-specific signatures in CP

Exocrine pancreatic tissues collected from the first cohort, organ donors (n=3) and CP (hereditary, n=5 and idiopathic n=4) patients, who underwent total pancreatectomy (table 1, online supplemental table 1,2 and online supplemental figure 1A), were used to isolate immune cells by immune cell enrichment method and fluorescence-activated cell sorting (FACS)^21^. Next, sorted live CD45^+^ pancreatic immune cells were further stained with 13 different surface protein antibodies for CITE-seq. The stained cells went through a droplet-based gel-bead barcoding system (10x genomics)^23^, which enables constructing independent maps of immunophenotypes of surface protein expressions, transcriptomes (5’ scRNA-seq), and the TCR repertoire (scTCR-seq) simultaneously in the same cells (figure 1A and online supplemental table 3). After quality control (online supplemental figure 1B) and removing doublets, we retained a total of 28,547 single cells and performed clustering analyses with gene expression data. This revealed 17 different cell populations that were visualized as uniform manifold approximation and projection (UMAP) embeddings (figure 1B and online supplemental data 1). Classification of cell populations was annotated by specific gene expressions that were inferred from various human single immune cell RNA-seq studies (figure 1C and online supplemental figure 2A)^24-26^. Major pancreatic immune cells that we identified from controls and CP were T cells, myeloid cells, B cells, and mast cells (figure 1B and online supplemental figure 2B). Noticeably, cells did not topologically cluster by experimental batch or individual subject, but cells from each group including control, hereditary, or idiopathic CP grouped together in a distinctive manner, highlighting the impact of disease and its etiology on the immune transcriptome (figure 1D and online supplemental figure 2C).

**Table 1.** Statistical comparisons of demographic and characteristics between control and CP subjects, a cohort for single-cell sequencing analysis.

| Characteristics | Control<br>(n=3) | CP<br>(n=9) | P value |
| --- | --- | --- | --- |
| Age (years, mean ± SD)<sup>A</sup> | 44.3 ± 8.2 | 26.2 ± 14.5 | 0.08 |
| Sex, m/f<sup>B</sup> | 1/2 | 3/6 | 1 |
| Height (meters, mean ± SD)<sup>A</sup> | 1.70 ± 0.41 | 1.62 ± 0.13 | 0.31 |
| Weight (kg, mean ± SD)<sup>A</sup> | 76.1 ± 18.5 | 61.6 ± 15.9 | 0.24 |
| Body mass index (kg/m<sup>2</sup>, mean ± SD)<sup>A</sup> | 26.1 ± 5.1 | 23.2 ± 4.1 | 0.37 |
| HbA1c (% , mean ±SD)<sup>A</sup> | 5.1 ± 0.1 | 5.2 ± 0.5 | 0.77 |
| SD, standard deviation; IQR, interquartile range. |  |  |  |
| <sup>A</sup>Unpaired two-tailed t-test; <sup>B</sup>Chi-square was used for p-values, Bold p value, p<0.05. |  |  |  |

**Figure 1.**
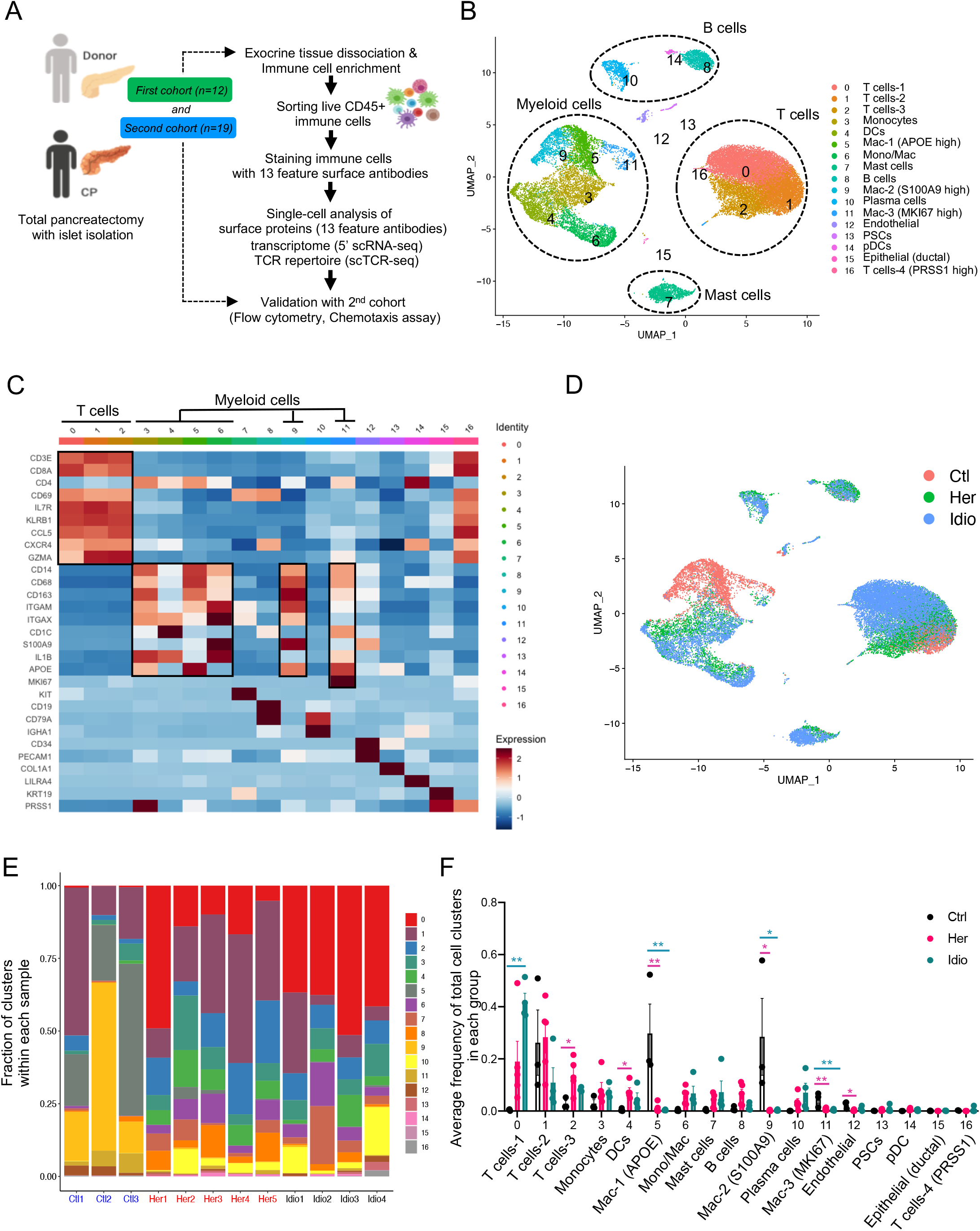
Human pancreatic immune cell transcriptional atlas. (**A**) Experimental design for single-cell sequencing (CITE-seq and TCR-seq) of pancreatic immune cells from CP patients and non-diseased control donors. (**B**) UMAP plot of all 28,547 pancreatic immune cells from CP patients (5 hereditary and 4 idiopathic CP) and 3 control donors presenting 17 clusters (Mac, macrophages; Mono/Mac, monocytes and macrophages; pDCs, plasmacytoid dendritic cells). (**C**) Heatmap of signature gene expression z scores across cells. (**D**) UMAP plot of pancreatic immune cells including 4,169 cells from control donors, 11,786 cells from hereditary CP, and 12,592 cells from idiopathic CP patients colored based on the group (Ctl, control; Her, hereditary CP; Idio, idiopathic CP). (**E**) Cell cluster frequency shown as a fraction of clusters from total cells in each patient or donor. (**F**) The frequency of 17 immune clusters defined by scRNA-seq clustering analysis shown as an average proportion of each cluster within each group. One-way ANOVA Kruskal-Wallis test (*p<0.05, **p<0.01, ***p<0.001). The comparison of each pair was differentiated by color: between Ctl and Her, Pink; between Ctl and Idio, Teal; between Her and Idio, Black.

We further analyzed the frequency and composition of immune populations within each group or individual subject by identified immune clusters. We found major immune cells that consist of control pancreas tissues were myeloid cells while a higher proportion of T cells contributed to pancreatic immune cells from CP tissues (figure 1E and online supplemental figure 3A, B). Among T cells and myeloid cells, distinct cell clusters were enriched in each group; macrophages including cluster #5, 9, and 11 were enriched in control, and distinct T cell clusters were expanded in CP groups. Cluster #2 was mainly enriched in hereditary CP while cluster #0 was predominant in idiopathic CP (figure 1F and online supplemental figure 3C, D). We also confirmed surface protein marker expression patterns in pancreatic immune cells such as CD3, CD8, CD11B, and HLA-DR by visualizing their expressions in the UMAP created by transcriptome data across all immune clusters (online supplemental figure 4A, B).

Next, we examined disease-specific gene signatures by identifying differentially expressed genes (DEGs) in CP versus control (online supplemental figure 5A, B). The top 20 upregulated genes in CP compared with controls included *HSP90AA1, FTH1, TNFAIP3, NFKBIA, NR4A1, CD69*, and *CCL20*. Furthermore, we performed Gene set enrichment analysis (GSEA) of DEGs in CP versus controls, and the results highlighted strong signatures for inflammatory responses including apoptosis, hypoxia, IL2-STAT5, interferon-gamma, and TNFA signaling (online supplemental figure 5C, D). Overall, single-cell transcriptome data analysis with pancreatic immune cells from control and CP tissues indicate disease-specific immune responses in local pathogenic area and distinct immune transcriptomic and protein expression signatures between hereditary and idiopathic CP.

### Distinct transcriptomic characteristics of pancreatic T cells between hereditary and idiopathic CP

One of the major immune populations infiltrating the pancreas of control and CP subjects were T cells. In order to scrutinize T cell transcriptomes in a higher resolution, we analyzed T cell populations (clusters # 0, 1, and 2 of total cell clusters, figure 1B) separately. We complied gene expression data from 15,913 cells for T cell clustering analyses and identified 13 different T cell clusters with population nomenclature annotated by specific marker gene expressions (figure 2A, B and online supplemental figure 6A). UMAP with T cell populations revealed markedly separated cell clusters by three groups, control, hereditary and idiopathic CP (figure 2C). Control cells comprised predominantly GZMA^+^ cytotoxic CD8^+^ T cells while distinct CD4^+^ or CD8^+^ T cell subpopulations constituted CP T cells (figure 2D, E and online supplemental figure 6B, C). Hereditary CP cells mainly consisted of CD4^+^ helper T cell (Th) subpopulations including CCR6^+^, TNF^+^ (Th1), regulatory T (Treg) cells, and HLA-DA^+^ CD8^+^ T cells. However, FTH1^+^ CD4^+^ or CD8^+^ T cells were predominant T cells in idiopathic CP (online supplemental figure 6D, E).

**Figure 2.**
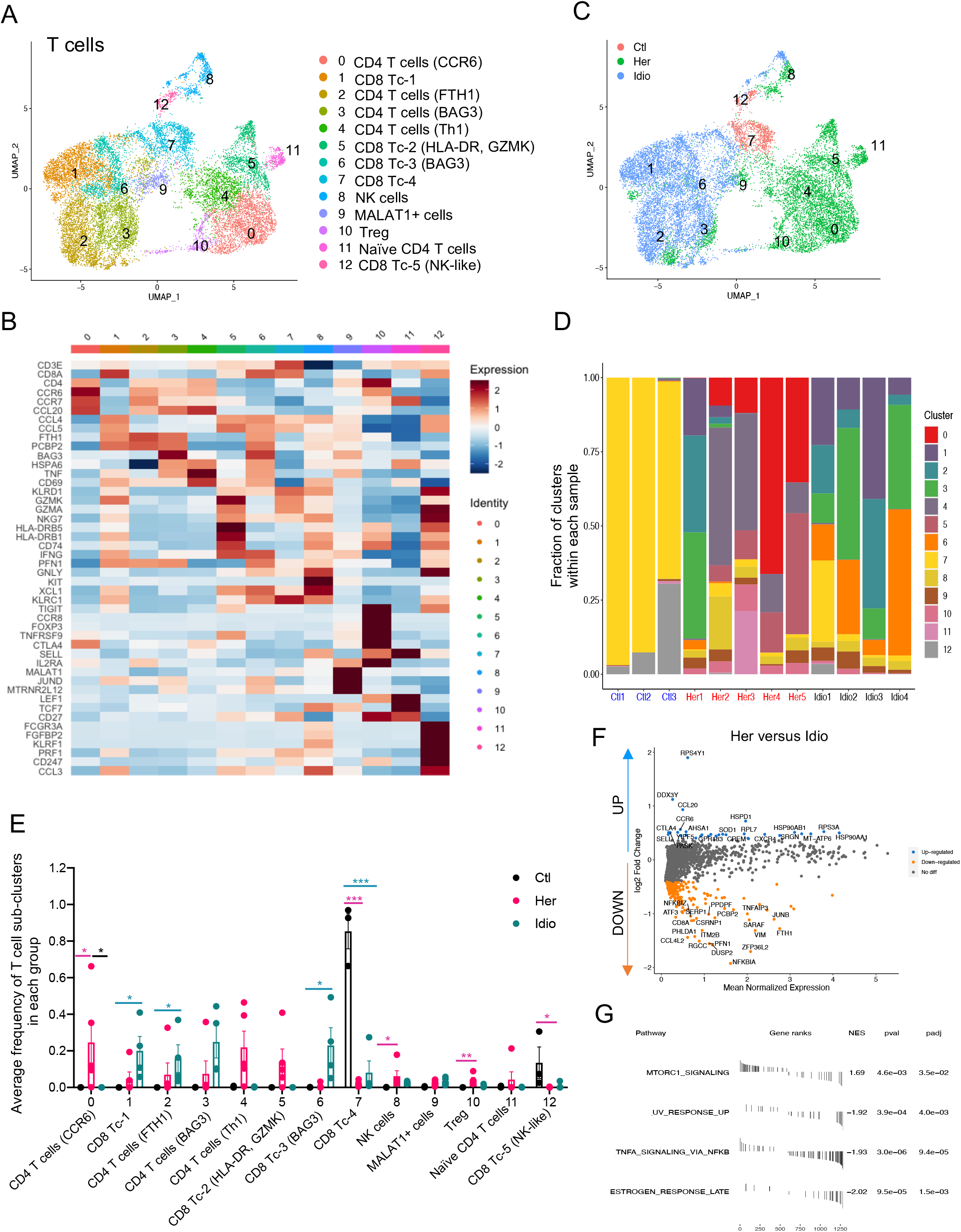
Distinct pancreatic T cell transcriptional signatures in hereditary versus idiopathic CP. (**A**) UMAP plot of 15,913 pancreatic T cells across controls and CP displaying 13 clusters. (**B**) Heatmap of signature gene expression z scores across cells. (**C**) UMAP plot of pancreatic T cells including 1,012 cells from control donors, 7,271 cells from hereditary CP, and 7,630 cells from idiopathic CP patients colored based on the group (Ctl, control; Her, hereditary CP; Idio, idiopathic CP). (**D**) Cell cluster frequency shown as a fraction of 13 T cell subclusters in each patient or donor. (**E**) The frequency of 13 T cell subclusters defined by scRNA-seq clustering analysis shown as an average proportion of each cluster within each group. One-way ANOVA Kruskal-Wallis test (*p<0.05, **p<0.01, ***p<0.001). The comparison of each pair was differentiated by color; between Ctl and Her, Pink; between Ctl and Idio, Teal; between Her and Idio, Black. (**F**) DEGs in between hereditary CP and idiopathic CP. Each dot represents a gene, with significantly up-regulated top 20 genes and down-regulated top 20 genes in hereditary CP versus idiopathic CP colored blue and yellow, respectively. (**F**) Functional enrichment analysis of significant hallmark gene sets in total T cells from hereditary CP versus idiopathic CP. NES, normalized enrichment score.

Furthermore, within these T cell subpopulations, we identified DEGs and performed GSEA of upregulated genes in CP versus control (online supplemental figure 7A, B), and in hereditary CP versus idiopathic CP (figure 2F, G). Some of the noticeable genes that were upregulated in hereditary compared with idiopathic CP included chemotactic receptors and ligands such as, *CCR6, CXCR4, GPR183*, and *CCL20*, suggesting their involvement in CD4^+^ T cell recruitment in hereditary CP.

### CD8^+^ T cell dependent unique TCR repertoire changes in CP

Next, we examined the TCR repertoire of the same T cells from which we extracted transcriptome data in control and CP. Paired transcripts, TCR alpha (TRA) and TCR beta (TRB), two distally encoded but co-expressed in single cells were sequenced and filtered through a programmed filtering system (Cell Ranger, 10x genomics). After barcode correction and trimming, V(D)J genes in the complementary determining region 3 (CDR3) of TCR transcripts were annotated. We identified a total of 12,856 unique paired ab TCR sequences and 5,180 T cells with paired TRA and TRB CDR3 sequences from the T cell repertoire of 12 control and CP subjects. In control T cells, 15.6 ± 6.0% of unique clonotypes were shared by 2 or more cells, which was significantly higher than that of hereditary CP T cells (4.03 ± 1.6%) (figure 3A). Corresponding to this result, the Gini-coefficient (an index of clonality) of control T cells was also significantly higher compared with the hereditary CP group indicating lower clonal expansion in hereditary CP T cells (figure 3B). Next, we assigned TCR sequences to cells with cluster identities as shown in Fig. 2A. Gini-coefficients of different T cell clusters revealed CD8^+^ T cell-skewed clonal expansion in control and CP groups (figure 3C). When we split T cells into CD8^+^ T cells and the rest, the Gini-coefficient of CD8^+^ T cells was significantly higher than that of the remainder T cells (figure 3D). This result was further confirmed by comparing UMAPs of *CD8A* gene expression and clonally expanded cell populations (figure 3E). Finally, we observed that Gini coefficients of subjects had a positive correlation with CD8^+^ T cell frequency in control and CP groups while Gini-coefficients had a negative correlation with CD4^+^ T cell frequency in all subjects (figure 3F). These data implicate that a significantly higher frequency of CD4^+^ T cells contributes to the lower Gini-coefficient and clonal expansion in hereditary CP compared with controls.

**Figure 3.**
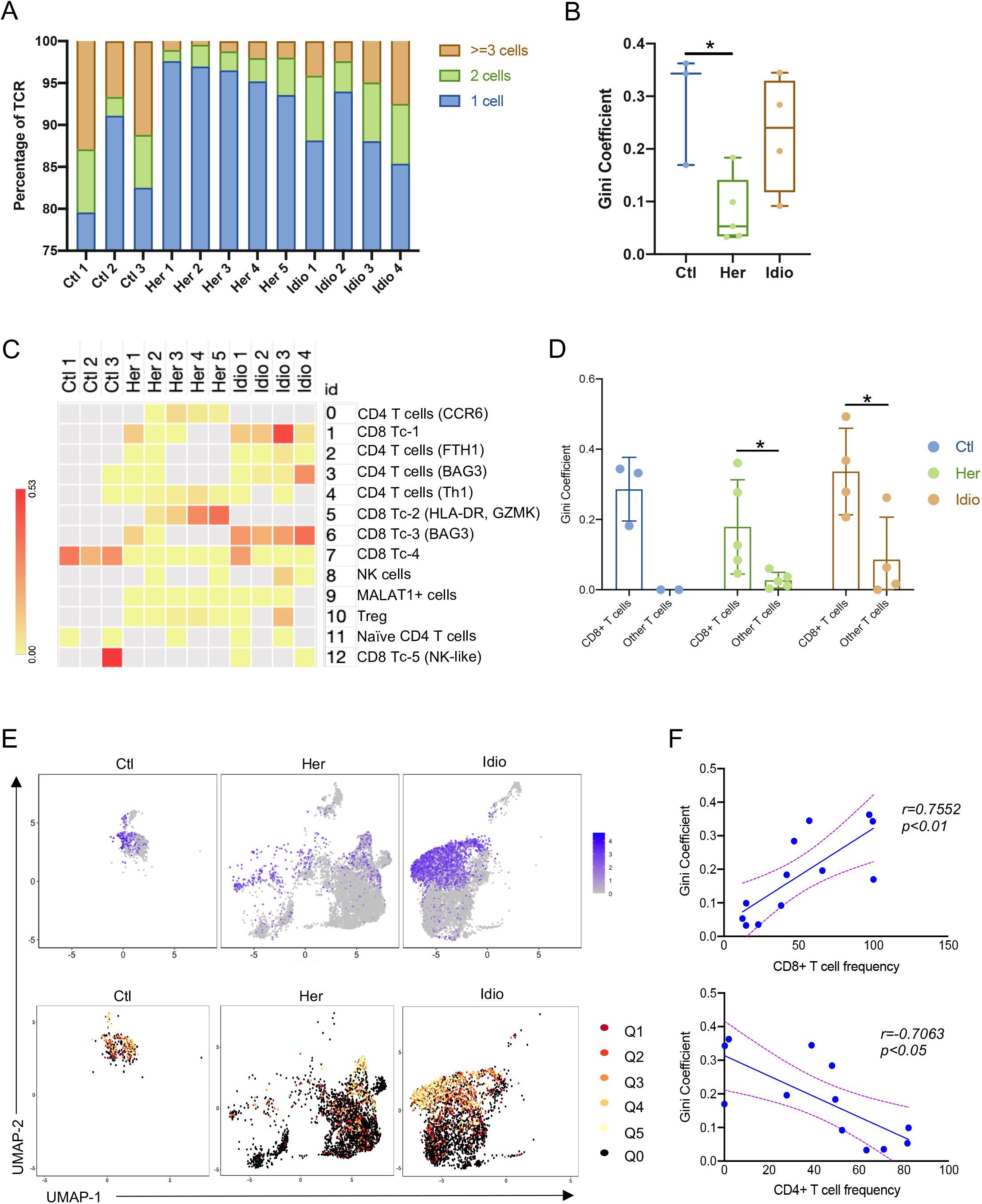
scRNA/TCR-seq unveils CD8^+^ T cell subset dependent unique TCR repertoire changes in CP. (**A**) Bar graph shows the percentage of unique paired TRA and TRB sequences that are shared by one cell (blue), by two cells (green), or three or more cells (orange) in each donor or CP patient. (**B**) Gini coefficients of control and CP group across all pancreatic T cell populations. For each group, a box and whisker plot is shown with the median and all values from minimum to maximum. One-way ANOVA with Tukey’s multiple comparison test, *p<0.05. (**C**) Heatmap showing Gini coefficients of pancreatic T cell clusters in each donor or patient. (**D**) Gini coefficients of pancreatic CD8^+^ T cells and other T cells in control and CP groups. (**E**) UMAP plot of *CD8A* gene expression in control or CP groups (top). TCR clonality on expression-driven UMAP overlay showing the distribution of clonal and unexpanded cell populations in cell clusters in control and CP groups (bottom). Q0, unexpanded; Q1, cells encompassing the top 20% of the most expanded clones in the group; Q2-4, cells representing the middle three quantiles; Q5, cells encompassing the bottom 20% of the most expanded clones. (**F**) Correlation analysis of Gini coefficients for pancreatic T cells and the frequency of CD8^+^ T cells (top) or CD4^+^ T cells (bottom) in control donors and CP patients.

These scRNA/TCR-seq results revealed that the degree of T cell clonal expansion or diversity in CP is mainly altered by the frequency of cytotoxic CD8^+^ T cells and infiltration of pathogenic CD4^+^ T cells. In hereditary CP, infiltrating CD4^+^ T cells likely dilute the effect of tissue-resident cytotoxic CD8^+^ T cells on TCR clonal expansion observed in control pancreatic tissues. This may provide key insight on CD4^+^ T cell-mediated pathogenic response especially in the case of hereditary CP.

### Unique interactions among T cell lineages and shared antigen-binding motifs in CP

In order to understand the connection and origin of T cell clusters, we examined the unique clonotypes (matched single-cell TCR-αβ profiling) and their overlaps between different clusters by analyzing clonotypes across all groups (figure 4A) or each group (online supplemental figure 8A, B, and C). Although higher Gini-coefficient and expanded clones were observed in control versus CP, no single clonotype was shared between different clusters in controls. Major clonotype overlaps were found in both hereditary and idiopathic CP (online supplemental figure 8A, B, and C). The greatest clonotype overlaps were found between CD8^+^ cytotoxic T cell clusters (Tc-1 and Tc-3) with 40 shared clonotypes, which were contributed by idiopathic CP. In fact, 63 unique shared clonotypes were found among CD8^+^ cytotoxic T cell clusters in idiopathic CP (figure 4A and online supplemental figure 8C). The next highest clonotype overlaps were found between CCR6^+^ CD4^+^ T cells and Th1 cells with 23 shared clonotypes, and these overlaps mainly occurred in hereditary CP (figure 4A and online supplemental figure 8B). These shared TCR clonotypes elucidated unique interactions among different T cell subtypes and clonal dynamics in each CP group. Specifically, a large number of shared clonotypes between CCR6^+^ CD4^+^ T cells and Th1 cells in hereditary CP suggests that these two Th cell subsets may have a shared origin. Interestingly, our analysis demonstrated overlaps between CD4^+^ and CD8^+^ T cell clusters, which may require further analyses regarding the length of CDR3 sequences in overlapped clonotypes as there is a higher probability that significantly short CDR3 sequences to be less impactful in recognizing a particular MHC class^27^.

**Figure 4.**
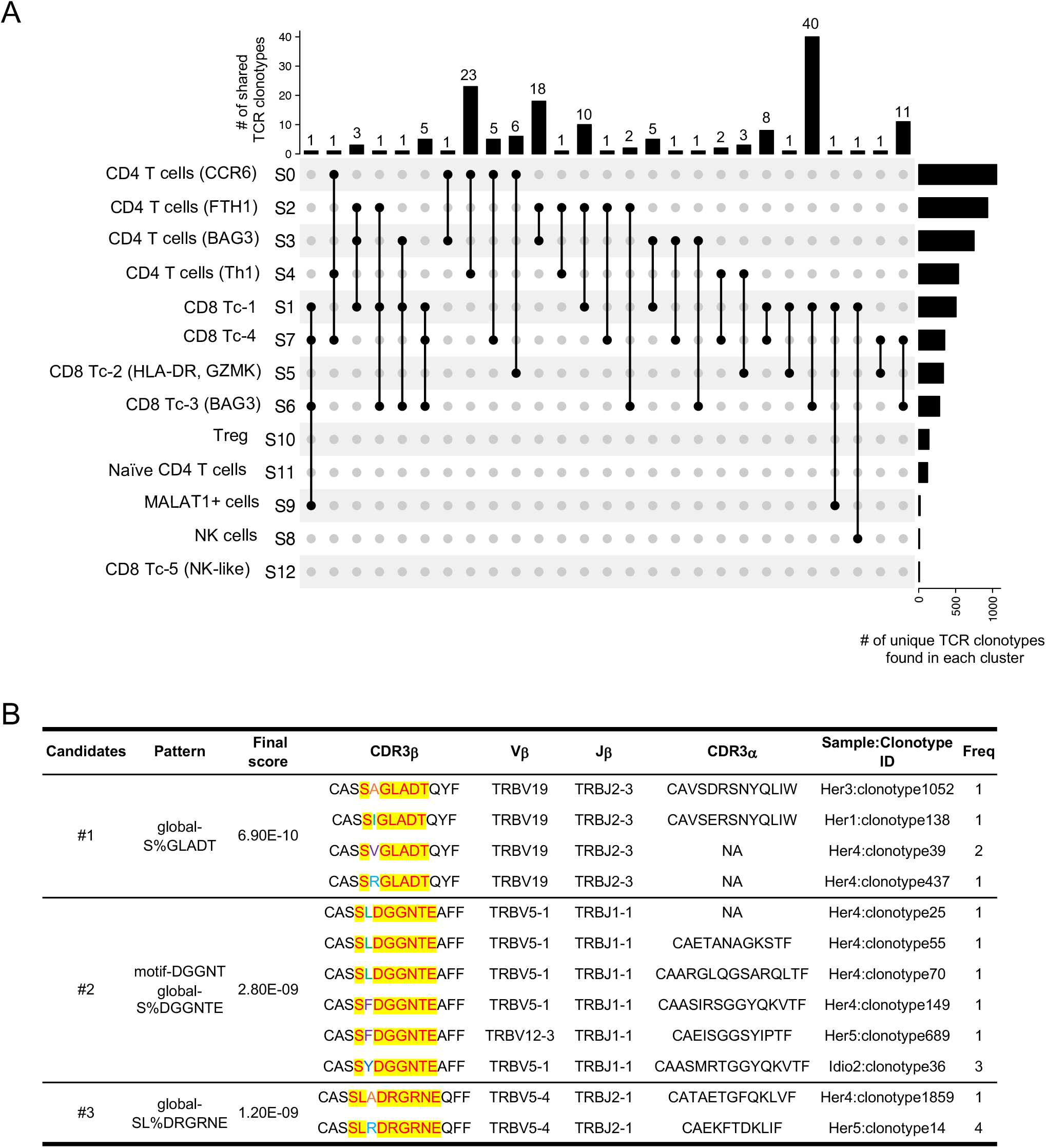
scRNA/TCR-seq uncovers unique interactions in T cell lineages and shared antigen binding motifs in CP. (**A**) Upset plots displaying TCR clonotypes shared among T cell clusters from T cells across control and CP groups. Each shared clonotype was indicated by black dots with a connected black line. The horizontal bar graph indicates the total number of shared TCR clonotypes for cluster intersections, and the vertical bar graph indicates the number of unique clonotypes found in a single cluster. (**B**) Representative TCR specificity groups and potential antigen binding motifs by GLIPH2 cluster analysis of TCR CDR3b sequences from pancreatic T cells across control, hereditary and idiopathic CP. Candidates were selected from the clusters with a final score of less than 10^−8^ and shared by at least two different individuals.

To examine whether pancreatic TCRs found in controls and CP recognize the same antigens, we performed GLIPH2 analysis which enables clustering TCRs that recognize the same epitope by screening shared antigen-binding motifs on the TCR CDR3β amino acid sequences^28^. A number of motifs shared by at least two different individuals from control donors and CP patients were selected based on final scoring, and among those, three clusters were shown as they also possess a certain level of similarity on their paired CDR3α sequences (figure 4B). The results showed that selected clusters were mainly shared by individuals from hereditary and idiopathic CP, not from controls. Remarkably, most of the TCRs from the three candidate clusters with unique common motifs were mainly shared by individuals from hereditary CP, which indicates T cells in hereditary CP have a higher possibility of reacting against common antigens or epitopes. Consistent with scRNA-seq results of T cells, this scRNA/TCR-seq data also supports that there are distinct immune responses, specifically T cell antigen reactions, between hereditary and idiopathic CP.

### Distinct transcriptional alteration of human pancreatic myeloid cells in CP

Myeloid cells have been known to play significant roles in CP and interact with non-immune cells such as PSCs^17^. Myeloid cells were another key contributor among pancreatic immune subsets found in the pancreas of control and CP subjects (figure 1B). When we further analyzed myeloid clusters separately (clusters # 3, 4, 5, 6, 9, and 11 in figure 1B), 11 different myeloid subclusters were identified that included monocytes (Mono), macrophages (Mac), and dendritic cells (DC) (figure 5A). Remarkably, cells belonging to each group (control, hereditary CP, or idiopathic CP) gathered together and topologically separated from each other with minimal overlaps when visualized in UMAP (figure 5B). Myeloid cell clusters were manually annotated by distinct cell marker gene expressions (figure 5C and online supplemental figure 9A). Importantly, the frequency of CD68^+^ and CD163^+^ macrophages and S100A8^+^ monocytes (clusters #3, 8) were significantly higher in controls compared with CP (figure 5D, E and online supplemental figure 9B, C). In CP, distinct monocyte and DC populations contributed to the myeloid compartment of hereditary or idiopathic CP immune cells. The most noticeable finding was a significantly higher frequency of CCL20^+^ monocytes in hereditary CP compared with control or idiopathic CP. Furthermore, DEG analysis in CP versus control revealed various inflammatory and chemoattractant molecules such as *IL1B, CXCL2, CXCL3, CXCL8*, and *CCL4*, which were significantly upregulated with enriched inflammatory signaling pathways in CP myeloid cells (online supplemental figure 9D, E). DEG analysis in hereditary versus idiopathic CP also identified various genes including *HLA-DR, CCL20*, and *IL1B* as significantly upregulated with enrichment of allograft rejection pathway in hereditary CP compared with idiopathic CP (figure 5F, G). This data further implicates that hereditary CP with higher HLA molecule expressions may associate with autoimmune responses although specific HLA typing would be necessary to confirm this notion^29 30^. Overall, single-cell transcriptional analysis of pancreatic myeloid populations revealed distinct myeloid subpopulation enrichment in each group; macrophages in controls, CCL20^+^ monocytes and IL1B^+^ DCs in hereditary CP, and FTH1^+^ or S100A8^+^ monocytes in idiopathic CP. Especially, significantly enriched CCL20^+^ monocyte population in hereditary CP suggests a potential role as a chemoattractant for the infiltrating CCR6^+^ CD4^+^ T cells.

**Figure 5.**
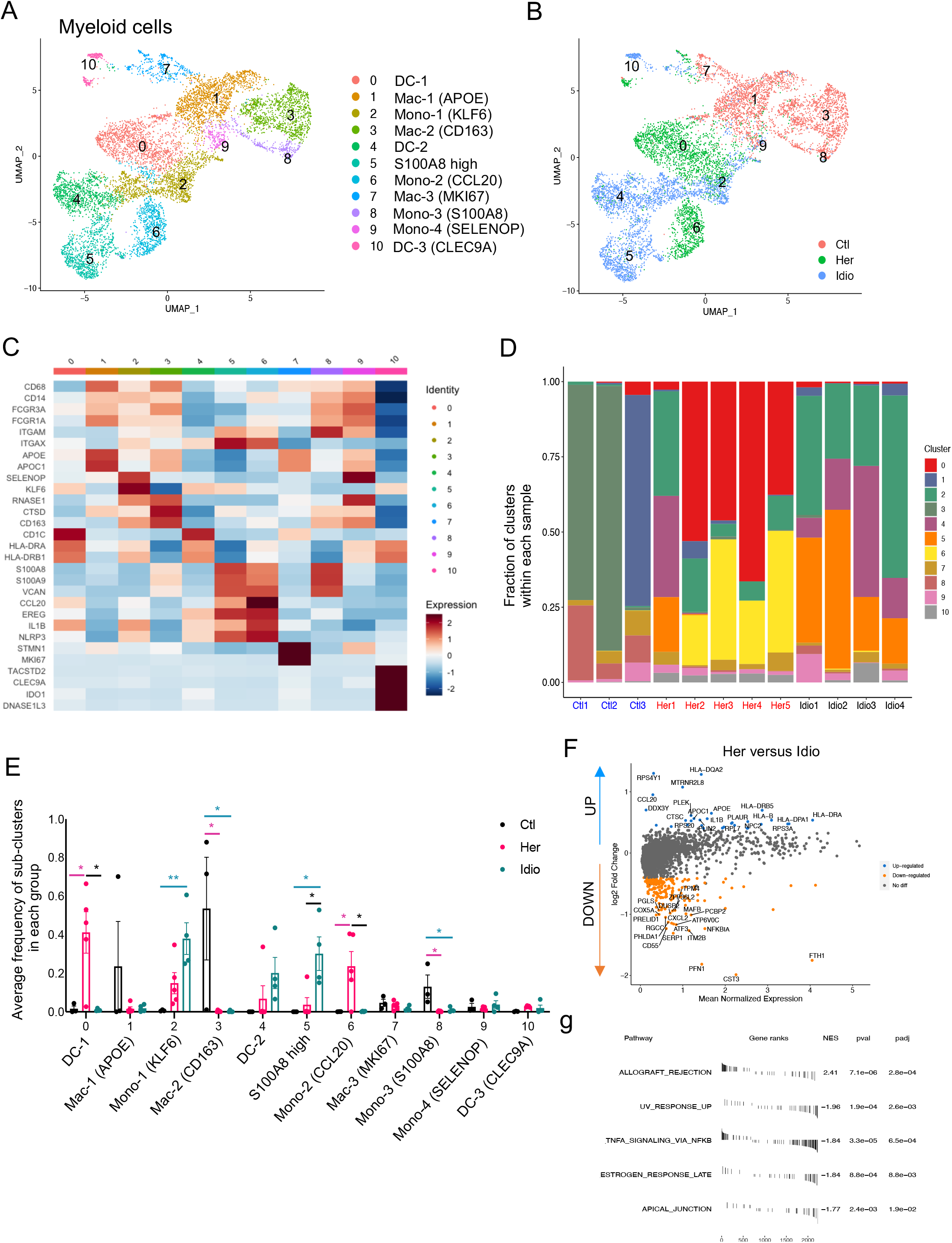
Distinct pancreatic myeloid cell transcriptional signatures between hereditary and idiopathic CP. (**A**) UMAP plot of 8,590 pancreatic myeloid cells presenting 11 clusters (DC, dendritic cells; Mac, macrophages; Mono, monocytes). (**B**) UMAP plot of pancreatic myeloid cells including 2,998 cells from control donors, 2,769 cells from hereditary CP, and 2,823 cells from idiopathic CP patients colored based on group (Ctl, control; Her, hereditary CP; Idio, idiopathic CP). (**C**) Heatmap of signature gene expression z scores across cells. (**D**) Cell cluster frequency shown as a fraction of 11 myeloid subclusters in each patient or donor. (**E**) The frequency of 11 myeloid subclusters shown as an average frequency of each cluster within each group. One-way ANOVA Kruskal-Wallis test (*p<0.05, **p<0.01). The comparison of each pair was differentiated by color; between Ctl and Her, Pink; between Ctl and Idio, Teal; between Her and Idio, Black. (**F**) DEGs between hereditary and idiopathic CP groups. Each dot represents a gene, with significantly up-regulated top 20 genes and down-regulated top 20 genes in hereditary CP versus idiopathic CP colored blue and yellow, respectively. (**G**) Functional enrichment analysis of significant hallmark gene sets comparing myeloid cells from hereditary CP with idiopathic CP. NES, normalized enrichment score.

### Functional analysis implicates the CCR6-CCL20 axis in hereditary CP

Consistent with the transcriptional expression of CCR6-CCL20 axis in CD4^+^ T cell and monocyte populations from hereditary CP, we confirmed this unique chemokine and receptor axis again by analyzing DEGs between hereditary and idiopathic CP within total immune populations (figure 6A). Accordingly, *CCR6* and *CCL20* were listed on the top 20 genes significantly upregulated in hereditary CP compared to idiopathic CP. Furthermore, to assess the crosstalk between myeloid cells and T cells, we analyzed cytokine or chemokine receptor-ligand interactions between myeloid cells and T cell compartment by combined expression of receptor-ligand (figure 6B). CCR6-CCL20 axis was uniquely expressed only in both CP groups, and hereditary CP had a higher combined expression of CCR6-CCL20 than idiopathic CP. Finally, we assessed the protein expression of CCR6 in T cells isolated from the second cohort of controls and CP subjects by flow cytometry (figure 6C, table 2, and online supplemental table 4,5). First, we were able to confirm a significantly reduced frequency of CD8^+^ T cells and a significantly higher frequency of CD4^+^ T cells in hereditary CP compared with controls.

**Figure 6.**
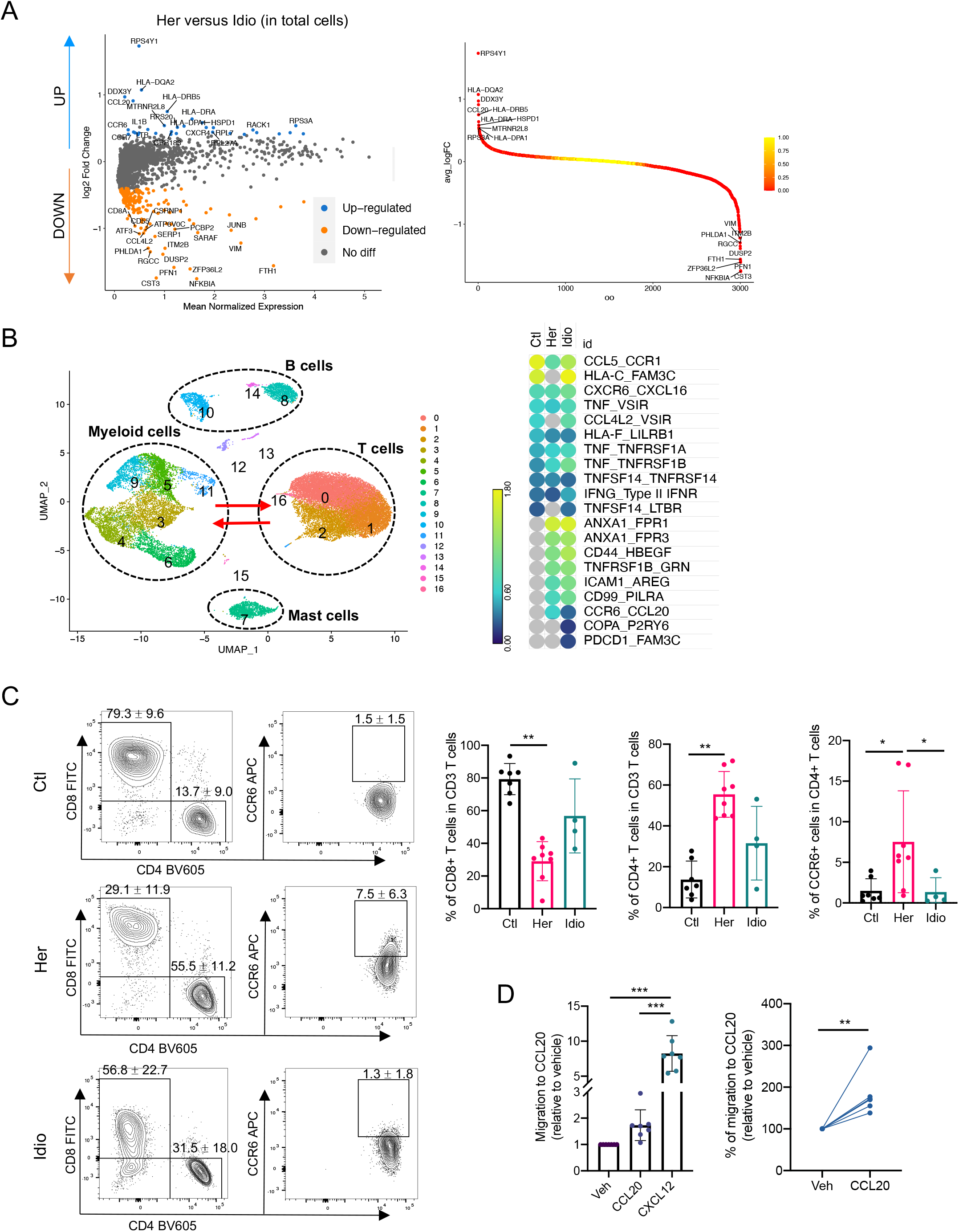
Functional analysis implicates CCR6-CCL20 axis as hereditary CP-specific pancreatic immune crosstalk. (**A**) Volcano plot of DEG analysis between hereditary and idiopathic CP groups with total immune populations. Each dot represents a gene, with significantly up-regulated top 20 genes and down-regulated top 20 genes in hereditary CP versus idiopathic CP colored blue and yellow, respectively (left). DEGs ranked by fold changes in hereditary CP versus idiopathic CP (right). (**B**) UMAP plot of total pancreatic immune cells (left). Heatmap of combined expression of selected cytokine/chemokine and their receptor displaying the interactions between myeloid cells and T cells discovered by CellphoneDB (right). (**C**) Flow cytometry analysis of CD8^+^, CD4^+^, or CCR6^+^ CD4^+^ T cells (Kruskal-Wallis test with Dunn’s multiple comparisons test, *p<0.05, **p<0.01). (**D**) Chemotaxis assay with pancreatic immune cells from hereditary CP (left, One-way ANOVA with Tukey’s multiple comparisons test; right, paired t-test; **p<0.01, ***p<0.001).

**Table 2.** Statistical comparisons of demographic and characteristics between control and CP subjects, a cohort for functional validation analyses.

| **Characteristics** | **Control**<br>**(n=7)** | **CP**<br>**(n=12)** | **P value** |
| --- | --- | --- | --- |
| Age (years, mean ± SD)<sup>A</sup> | 31.1 ± 9.6 | 27.5 ± 12.3 | 0.51 |
| Sex, m/f<sup>B</sup> | 4/3 | 6/6 | 0.76 |
| Height (meters, mean ± SD)<sup>A</sup> | 1.69 ± 0.41 | 1.67 ± 0.90 | 0.58 |
| Weight (kg, mean ± SD)<sup>A</sup> | 84.8 ± 11.2 | 70.2 ± 19.7 | 0.09 |
| Body mass index (kg/m<sup>2</sup>, mean ± SD)<sup>A</sup> | 29.6 ± 3.5 | 24.8 ± 5.3 | 0.05 |
| HbA1c (% , mean ±SD)<sup>A</sup> | 5.3 ± 0.6 | 5.3 ± 0.3 | 0.97 |
| SD, standard deviation; IQR, interquartile range. |  |  |  |
| <sup>A</sup>Unpaired two-tailed t-test; <sup>B</sup>Chi-square was used for p-values, Bold p value, p<0.05. |  |  |  |

Consistent with scRNA-seq results, the frequency of CD8^+^ or CD4^+^ T cells in idiopathic CP was in the middle, ranging between that of controls and hereditary CP. More importantly, the percentage of CCR6 expression in CD4^+^ T cells was significantly augmented in hereditary CP compared with controls or idiopathic CP. After we confirmed the elevated protein expression of CCR6 in CD4^+^ T cells in hereditary CP, we next assessed the functional significance of the CCR6 expression in T cells by CCL20-mediated cell migration assay (figure 6D). Transwell chemotaxis was performed, and CXCL12 was used as a strong positive control, which is standard in T cell migration assays^31^. Pancreatic immune cells from hereditary CP tissues migrated to soluble, recombinant human CCL20 at a concentration comparable to previous reports^32 33^. The increased chemotactic responsiveness of the pancreas infiltrating T cells to CCL20 confirmed the functional significance of the increased receptor (CCR6) expression in hereditary CP. These data suggest that T cells migrate and infiltrate the diseased pancreas in hereditary CP through the CCR6-CCL20 axis.

## DISCUSSION

Human CP studies have been hindered in part due to limited access to clinical specimens. By leveraging the ability to access large fractions of exocrine pancreas tissues from organ donors and CP patients undergoing islet isolation, we were able to perform in-depth single-cell level immune analyses using state of the art technologies. Our studies did not include CP related to alcohol due to an insufficient number of patients undergoing total pancreatectomy with islet transplantation with this etiology^34^, but we were able to receive a sufficient number of cases with hereditary and idiopathic CP for in-depth immune analyses. Our previous study using flow cytometry and bulk TCR-sequencing revealed distinct immune characteristics between hereditary and idiopathic CP highlighting unprecedented critical insights into distinctive disease pathogenesis of CP with different etiologies^21^. To further delineate pathogenic signals and uncover differences in immune responses underlying different etiologies of CP, here we used integrative single-cell multi-omics sequencing analyses to assess the TCR repertoire and RNA/protein expressions simultaneously at a single-cell level. These unbiased in-depth analytic technologies enabled comprehensive comparison between control and CP, as well as between different CP groups and identified novel immune subsets, their crosstalk, and T cell subset based TCR repertoire changes. Consistent with our previous study^21^, hereditary and idiopathic CP had distinct immune characteristics not only in frequencies of immune subpopulations but also in their unique gene expression patterns and the degree of T cell clonal expansions among different subsets. Furthermore, T cell analyses revealed a significant portion of CD4^+^ T cell subsets (*CCD6*^*+*^ Th, *TNF*^*+*^ Th1, and Treg) were enriched in the hereditary CP pancreas highlighting the immune signature of hereditary CP pancreas is largely conserved across heterogeneous CD4^+^ T cell subsets. However, idiopathic CP showed evenly distributed CD4^+^ and CD8^+^ T cell subsets uniquely with *FTH1* and *PCBP2* expressions, which indicate that human idiopathic CP is highly associated with iron metabolism in immune cells and dysregulated iron metabolism has been reported in pancreatitis^35-37^.

Distinct T cell characteristics between hereditary and idiopathic CP were also seen in the TCR repertoire analysis. Pancreatic T cells revealed an increased clonal expansion in CD8^+^ T cell subsets in all control and CP groups, which led to significantly lower T cell clonality in hereditary CP where CD4^+^ T cell infiltrates are increased. These data suggest that differences in the TCR repertoire found in control versus CP or hereditary versus idiopathic CP were highly relevant to the distribution of specific T cell subsets. In contrast to reports of TCR repertoires in auto-immune, infectious, and malignant diseases^24 38-40^ where newly infiltrating immune cells are usually responsible for the increased clonality, here infiltrating pancreatic CD4^+^ T cells contribute to the reduced clonality in the diseased states but increased heterogeneity of overall T cell subsets. In addition, shared TCR clone analysis with scRNA/TCR-seq data unveiled the unique interactions between CCR6^+^ Th and Th1 subsets in hereditary CP supporting naturally increased CD4^+^ T cell heterogeneity in the microenvironment of hereditary CP. The unique pancreas-specific TCR repertoire found in hereditary and idiopathic CP will contribute to a better understanding of pathogenic T cell infiltration and behavior, which have not been reported previously and likely to provide fundamental clues of pathogenic mechanisms and/or disease progression in CP. Given distinct TCR repertoire changes and common antigen-binding motifs found in hereditary versus idiopathic CP, future studies that include predicting HLA restriction and antigen screening against disease-specific TCR candidates^41^ will be important.

One of the striking findings in the single-cell analyses was the upregulation of the CCR6-CCL20 axis in hereditary CP. Upregulation of CCL20 by pro-inflammatory cytokines such as IL-1b and TNF-a^42-44^ and infiltrating CCR6^+^ lymphocytes have been found in microenvironments of inflammatory, infectious, and malignant states in various organs such as the gut, intestine, liver, and lung^32 45-48^. CCL20 expression pattern has been compared in pancreatic cancer versus CP^49^ and its tumor-promoting role in pancreatic cancer has been proposed^50 51^, however, CCR6^+^ T cell infiltration in hereditary or any type of CP to our knowledge has not been reported. Our findings also support CP-specific crosstalk between CD4^+^ T cells and monocytes through the CCR6-CCL20 axis in hereditary CP, a novel insight into possible precision-targeting of this disease. Interestingly, among the different causes of CP, hereditary CP has the highest risk for developing pancreatic cancer with a cumulative increased risk of up to 40%^52 53^. The significant upregulation of the CCL20-CCR6 axis in hereditary as compared to idiopathic CP (and organ donors) might contribute to the increased risk of pancreas cancer in hereditary CP although further studies have to be done to confirm this correlation.

We focused our analysis and reporting on the major immune cell subsets, but we also found that B cells and mast cells were distinctly distributed in hereditary versus idiopathic CP although their frequencies were limited to less than 10% of total immune cells. It will be important in the future to analyze these cells in-depth and examine their potential functional significance in CP since there are several reports suggesting their potential pathogenic roles in pancreatitis^54-56^. Collectively, our approaches with integrative single-cell analyses unveiled distinct pancreatic immune signatures and pathways between different etiologies of CP, thereby expanding our understanding of pancreatic immune cell signaling and function in the disease-specific pathogenic microenvironment. In addition, our study contributes to the growing literature of gaining insight into the characterization and function of human pancreas immune cells in health and disease.

## Acknowledgments

We thank our collaborators, patients, and donors who provided precious human pancreatic tissues for this study. We also thank Y. Yang, Y. Wei, and A.R. Ji for technical assistance, and all other members of the Habtezion laboratory for their helpful comments and suggestions.

## Funding

This study was supported in part by NIH grant DK105263 (AH), NPF Research Grant (BL), and Diabetes Genomics and Analysis Core of the Stanford Diabetes Research Center (P30DK116074).

## Author contributions

B.L. and A.H. designed experiments and wrote manuscript. A.H. provided overall guide and supervision. M.D.B. and D.H. provided CP patients’ tissues and clinical information. G.L.S provided control pancreatic tissues and clinical information. B.L. performed all experiments and analyzed data, and H.N. performed FACS to prepare single cells for single-cell sequencing, flow cytometry panel design, and data analysis. Y.Y. analyzed CITE-seq and scTCR-seq data. H.H. performed GLIPH2 analysis, interpreted and reviewed scTCR-seq results. M.M.D., S.J.P., and M.D.B reviewed manuscript and participated in the interpretation of data.

## Competing interests

Authors declare that they have no competing interests.

## Patient and public involvement

Patients or the public were not involved in the design, conduct, reporting, or dissemination plans of this research.

## Patient consent for publication

Not required.

## Ethics approval

This study was approved by Institutional Review Board of the Stanford University. Patients or the public were not involved in the design, conduct, reporting, or dissemination plans of this research.

## Data and materials availability

All raw and processed sequencing data have been deposited with Gene Expression Omnibus database, GSE165045. Code used for data analysis is available at https://github.com/yangysheep2018/CITE-seq-TCR_paper. All other processed data are available in the main text or the supplementary materials.

## License statement

This is an open access article distributed in accordance with the Creative Commons Attribution Non Commercial (CC BY-NC 4.0) license, which permits others to distribute, remix, adapt, build upon this work non-commercially, and license their derivative works on different terms, provided the original work is properly cited, appropriate credit is given, any changes made indicated, and the use is non-commercial.

## Notes

### Competing Interest Statement

The authors have declared no competing interest.

### Summary of Updates

ORCID number of Greg Szot is updated.

## REFERENCES

1. Witt H, Apte MV, Keim V, et al. Chronic pancreatitis: challenges and advances in pathogenesis, genetics, diagnosis, and therapy. Gastroenterology 2007;132(4):1557–73. doi: 10.1053/j.gastro.2007.03.001 [published Online First: 2007/05/01]

2. Yadav D, Timmons L, Benson JT, et al. Incidence, prevalence, and survival of chronic pancreatitis: a population-based study. Am J Gastroenterol 2011;106(12):2192–9. doi: 10.1038/ajg.2011.328 [published Online First: 2011/09/29]

3. Majumder S, Chari ST. Chronic pancreatitis. Lancet 2016;387(10031):1957–66. doi: 10.1016/s0140-6736(16)00097-0 [published Online First: 2016/03/08]

4. Kleeff J, Whitcomb DC, Shimosegawa T, et al. Chronic pancreatitis. Nat Rev Dis Primers 2017;3:17060. doi: 10.1038/nrdp.2017.60 [published Online First: 2017/09/08]

5. Whitcomb DC. Genetic aspects of pancreatitis. Annu Rev Med 2010;61:413–24. doi: 10.1146/annurev.med.041608.121416 [published Online First: 2010/01/12]

6. Muniraj T, Aslanian HR, Farrell J, et al. Chronic pancreatitis, a comprehensive review and update. Part I: epidemiology, etiology, risk factors, genetics, pathophysiology, and clinical features. Dis Mon 2014;60(12):530–50. doi: 10.1016/j.disamonth.2014.11.002 [published Online First: 2014/12/17]

7. Singh VK, Yadav D, Garg PK. Diagnosis and Management of Chronic Pancreatitis: A Review. JAMA 2019;322(24):2422–34. doi: 10.1001/jama.2019.19411 [published Online First: 2019/12/21]

8. Abu-El-Haija M, Nathan JD. Pediatric chronic pancreatitis: Updates in the 21st century. Pancreatology 2018;18(4):354–59. doi: 10.1016/j.pan.2018.04.013 [published Online First: 2018/05/05]

9. Aghdassi AA, Mayerle J, Christochowitz S, et al. Animal models for investigating chronic pancreatitis. Fibrogenesis Tissue Repair 2011;4(1):26. doi: 10.1186/1755-1536-4-26 [published Online First: 2011/12/03]

10. Klauss S, Schorn S, Teller S, et al. Genetically induced vs. classical animal models of chronic pancreatitis: a critical comparison. FASEB J 2018: fj201800241RR. doi: 10.1096/fj.201800241RR [published Online First: 2018/06/05]

11. Suda K, Takase M, Fukumura Y, et al. Histopathologic difference between chronic pancreatitis animal models and human chronic pancreatitis. Pancreas 2004;28(3):e86–9. doi: 10.1097/00006676-200404000-00030 [published Online First: 2004/04/16]

12. Pasca di Magliano M, Forsmark C, Freedman S, et al. Advances in acute and chronic pancreatitis: from development to inflammation and repair. Gastroenterology 2013;144(1):e1–4. doi: 10.1053/j.gastro.2012.11.018 [published Online First: 2012/11/20]

13. Habtezion A. Inflammation in acute and chronic pancreatitis. Curr Opin Gastroenterol 2015;31(5):395–9. doi: 10.1097/MOG.0000000000000195 [published Online First: 2015/06/25]

14. Watanabe T, Kudo M, Strober W. Immunopathogenesis of pancreatitis. Mucosal Immunol 2017;10(2):283–98. doi: 10.1038/mi.2016.101 [published Online First: 2016/11/17]

15. Xu S, Chheda C, Ouhaddi Y, et al. Characterization of Mouse Models of Early Pancreatic Lesions Induced by Alcohol and Chronic Pancreatitis. Pancreas 2015;44(6):882–7. doi: 10.1097/MPA.0000000000000380 [published Online First: 2015/07/15]

16. Xiao X, Fischbach S, Zhang T, et al. SMAD3/Stat3 Signaling Mediates beta-Cell Epithelial-Mesenchymal Transition in Chronic Pancreatitis-Related Diabetes. Diabetes 2017;66(10):2646–58. doi: 10.2337/db17-0537 [published Online First: 2017/08/05]

17. Xue J, Sharma V, Hsieh MH, et al. Alternatively activated macrophages promote pancreatic fibrosis in chronic pancreatitis. Nat Commun 2015;6:7158. doi: 10.1038/ncomms8158 [published Online First: 2015/05/20]

18. Xue J, Zhao Q, Sharma V, et al. Aryl Hydrocarbon Receptor Ligands in Cigarette Smoke Induce Production of Interleukin-22 to Promote Pancreatic Fibrosis in Models of Chronic Pancreatitis. Gastroenterology 2016;151(6):1206–17. doi: 10.1053/j.gastro.2016.09.064 [published Online First: 2016/11/05]

19. Bhatia R, Thompson C, Ganguly K, et al. Alcohol and Smoking Mediated Modulations in Adaptive Immunity in Pancreatitis. Cells 2020;9(8) doi: 10.3390/cells9081880 [published Online First: 2020/08/17]

20. Zhao Q, Manohar M, Wei Y, et al. STING signalling protects against chronic pancreatitis by modulating Th17 response. Gut 2019;68(10):1827–37. doi: 10.1136/gutjnl-2018-317098 [published Online First: 2019/02/02]

21. Lee B, Adamska JZ, Namkoong H, et al. Distinct immune characteristics distinguish hereditary and idiopathic chronic pancreatitis. J Clin Invest 2020;130(5):2705–11. doi: 10.1172/JCI134066 [published Online First: 2020/02/14]

22. Stoeckius M, Hafemeister C, Stephenson W, et al. Simultaneous epitope and transcriptome measurement in single cells. Nat Methods 2017;14(9):865–68. doi: 10.1038/nmeth.4380 [published Online First: 2017/08/02]

23. Zheng GX, Terry JM, Belgrader P, et al. Massively parallel digital transcriptional profiling of single cells. Nat Commun 2017;8:14049. doi: 10.1038/ncomms14049 [published Online First: 2017/01/17]

24. Oh DY, Kwek SS, Raju SS, et al. Intratumoral CD4(+) T Cells Mediate Anti-tumor Cytotoxicity in Human Bladder Cancer. Cell 2020;181(7):1612–25 e13. doi: 10.1016/j.cell.2020.05.017 [published Online First: 2020/06/05]

25. Zhao J, Zhang S, Liu Y, et al. Single-cell RNA sequencing reveals the heterogeneity of liver-resident immune cells in human. Cell Discov 2020;6:22. doi: 10.1038/s41421-020-0157-z [published Online First: 2020/05/01]

26. Luoma AM, Suo S, Williams HL, et al. Molecular Pathways of Colon Inflammation Induced by Cancer Immunotherapy. Cell 2020;182(3):655–71 e22. doi: 10.1016/j.cell.2020.06.001 [published Online First: 2020/07/01]

27. Carter JA, Preall JB, Grigaityte K, et al. Single T Cell Sequencing Demonstrates the Functional Role of alphabeta TCR Pairing in Cell Lineage and Antigen Specificity. Front Immunol 2019;10:1516. doi: 10.3389/fimmu.2019.01516 [published Online First: 2019/08/17]

28. Huang H, Wang C, Rubelt F, et al. Analyzing the Mycobacterium tuberculosis immune response by T-cell receptor clustering with GLIPH2 and genome-wide antigen screening. Nat Biotechnol 2020;38(10):1194–202. doi: 10.1038/s41587-020-0505-4 [published Online First: 2020/04/29]

29. Thomson G. HLA disease associations: models for the study of complex human genetic disorders. Crit Rev Clin Lab Sci 1995;32(2):183–219. doi: 10.3109/10408369509084684 [published Online First: 1995/01/01]

30. Caillat-Zucman S. Molecular mechanisms of HLA association with autoimmune diseases. Tissue Antigens 2009;73(1):1–8. doi: 10.1111/j.1399-0039.2008.01167.x [published Online First: 2008/11/20]

31. Bleul CC, Fuhlbrigge RC, Casasnovas JM, et al. A highly efficacious lymphocyte chemoattractant, stromal cell-derived factor 1 (SDF-1). J Exp Med 1996;184(3):1101–9. doi: 10.1084/jem.184.3.1101 [published Online First: 1996/09/01]

32. Wu YY, Tsai HF, Lin WC, et al. Upregulation of CCL20 and recruitment of CCR6+ gastric infiltrating lymphocytes in Helicobacter pylori gastritis. Infect Immun 2007;75(9):4357–63. doi: 10.1128/IAI.01660-06 [published Online First: 2007/06/15]

33. Francis JN, Sabroe I, Lloyd CM, et al. Elevated CCR6+ CD4+ T lymphocytes in tissue compared with blood and induction of CCL20 during the asthmatic late response. Clin Exp Immunol 2008;152(3):440–7. doi: 10.1111/j.1365-2249.2008.03657.x [published Online First: 2008/04/22]

34. Chinnakotla S, Beilman GJ, Dunn TB, et al. Factors Predicting Outcomes After a Total Pancreatectomy and Islet Autotransplantation Lessons Learned From Over 500 Cases. Ann Surg 2015;262(4):610–22. doi: 10.1097/SLA.0000000000001453 [published Online First: 2015/09/15]

35. Sledzinski M, Borkowska A, Sielicka-Dudzin A, et al. Cerulein-induced acute pancreatitis is associated with c-Jun NH(2)-terminal kinase 1-dependent ferritin degradation and iron-dependent free radicals formation. Pancreas 2013;42(7):1070–7. doi: 10.1097/MPA.0b013e318287d097 [published Online First: 2013/08/08]

36. Lunova M, Schwarz P, Nuraldeen R, et al. Hepcidin knockout mice spontaneously develop chronic pancreatitis owing to cytoplasmic iron overload in acinar cells. J Pathol 2017;241(1):104–14. doi: 10.1002/path.4822 [published Online First: 2016/10/16]

37. Chand SK, Singh RG, Pendharkar SA, et al. Interplay between innate immunity and iron metabolism after acute pancreatitis. Cytokine 2018;103:90–98. doi: 10.1016/j.cyto.2017.09.014 [published Online First: 2017/10/07]

38. Savas P, Virassamy B, Ye C, et al. Single-cell profiling of breast cancer T cells reveals a tissue-resident memory subset associated with improved prognosis. Nat Med 2018;24(7):986–93. doi: 10.1038/s41591-018-0078-7 [published Online First: 2018/06/27]

39. Allez M, Auzolle C, Ngollo M, et al. T cell clonal expansions in ileal Crohn’s disease are associated with smoking behaviour and postoperative recurrence. Gut 2019;68(11):1961–70. doi: 10.1136/gutjnl-2018-317878 [published Online First: 2019/02/23]

40. Gantner P, Pagliuzza A, Pardons M, et al. Single-cell TCR sequencing reveals phenotypically diverse clonally expanded cells harboring inducible HIV proviruses during ART. Nat Commun 2020;11(1):4089. doi: 10.1038/s41467-020-17898-8 [published Online First: 2020/08/17]

41. Gee MH, Han A, Lofgren SM, et al. Antigen Identification for Orphan T Cell Receptors Expressed on Tumor-Infiltrating Lymphocytes. Cell 2018;172(3):549–63 e16. doi: 10.1016/j.cell.2017.11.043 [published Online First: 2017/12/26]

42. Fujiie S, Hieshima K, Izawa D, et al. Proinflammatory cytokines induce liver and activation-regulated chemokine/macrophage inflammatory protein-3alpha/CCL20 in mucosal epithelial cells through NF-kappaB [correction of NK-kappaB]. Int Immunol 2001;13(10):1255–63. doi: 10.1093/intimm/13.10.1255 [published Online First: 2001/10/03]

43. Hirota K, Yoshitomi H, Hashimoto M, et al. Preferential recruitment of CCR6-expressing Th17 cells to inflamed joints via CCL20 in rheumatoid arthritis and its animal model. J Exp Med 2007;204(12):2803–12. doi: 10.1084/jem.20071397 [published Online First: 2007/11/21]

44. Iwamoto S, Kido M, Aoki N, et al. TNF-α is essential in the induction of fatal autoimmune hepatitis in mice through upregulation of hepatic CCL20 expression. Clin Immunol 2013;146(1):15–25. doi: 10.1016/j.clim.2012.10.008 [published Online First: 2012/11/28]

45. Facco M, Baesso I, Miorin M, et al. Expression and role of CCR6/CCL20 chemokine axis in pulmonary sarcoidosis. J Leukoc Biol 2007;82(4):946–55. doi: 10.1189/jlb.0307133 [published Online First: 2007/07/07]

46. Lee AY, Eri R, Lyons AB, et al. CC Chemokine Ligand 20 and Its Cognate Receptor CCR6 in Mucosal T Cell Immunology and Inflammatory Bowel Disease: Odd Couple or Axis of Evil? Front Immunol 2013;4:194. doi: 10.3389/fimmu.2013.00194 [published Online First: 2013/07/23]

47. Ranasinghe R, Eri R. Modulation of the CCR6-CCL20 Axis: A Potential Therapeutic Target in Inflammation and Cancer. Medicina (Kaunas) 2018;54(5) doi: 10.3390/medicina54050088 [published Online First: 2018/11/21]

48. Planas D, Zhang Y, Monteiro P, et al. HIV-1 selectively targets gut-homing CCR6+CD4+ T cells via mTOR-dependent mechanisms. JCI Insight 2017;2(15) doi: 10.1172/jci.insight.93230 [published Online First: 2017/08/05]

49. Klemm C, Dommisch H, Goke F, et al. Expression profiles for 14-3-3 zeta and CCL20 in pancreatic cancer and chronic pancreatitis. Pathol Res Pract 2014;210(6):335–41. doi: 10.1016/j.prp.2014.01.001 [published Online First: 2014/03/19]

50. Liu B, Jia Y, Ma J, et al. Tumor-associated macrophage-derived CCL20 enhances the growth and metastasis of pancreatic cancer. Acta Biochim Biophys Sin (Shanghai) 2016;48(12):1067–74. doi: 10.1093/abbs/gmw101 [published Online First: 2016/11/01]

51. Geismann C, Grohmann F, Dreher A, et al. Role of CCL20 mediated immune cell recruitment in NF-kappaB mediated TRAIL resistance of pancreatic cancer. Biochim Biophys Acta Mol Cell Res 2017;1864(5):782–96. doi: 10.1016/j.bbamcr.2017.02.005 [published Online First: 2017/02/12]

52. Lowenfels AB, Maisonneuve P, DiMagno EP, et al. Hereditary pancreatitis and the risk of pancreatic cancer. International Hereditary Pancreatitis Study Group. J Natl Cancer Inst 1997;89(6):442–6. doi: 10.1093/jnci/89.6.442 [published Online First: 1997/03/19]

53. Weiss FU. Pancreatic cancer risk in hereditary pancreatitis. Front Physiol 2014;5:70. doi: 10.3389/fphys.2014.00070 [published Online First: 2014/03/07]

54. Esposito I, Friess H, Kappeler A, et al. Mast cell distribution and activation in chronic pancreatitis. Hum Pathol 2001;32(11):1174–83. doi: 10.1053/hupa.2001.28947 [published Online First: 2001/12/01]

55. Demir IE, Schorn S, Schremmer-Danninger E, et al. Perineural mast cells are specifically enriched in pancreatic neuritis and neuropathic pain in pancreatic cancer and chronic pancreatitis. PLoS One 2013;8(3):e60529. doi: 10.1371/journal.pone.0060529 [published Online First: 2013/04/05]

56. Qiu L, Zhou Y, Yu Q, et al. Decreased levels of regulatory B cells in patients with acute pancreatitis: association with the severity of the disease. Oncotarget 2018;9(90):36067–82. doi: 10.18632/oncotarget.23911 [published Online First: 2018/12/14]

