## Supplemental Materials for "Single-cell sequencing unveils distinct immune microenvironment with CCR6-CCL20 crosstalk in human chronic pancreatitis"

**Supplemental Methods**

***Human pancreatic specimens***

Human pancreatic tissues from deceased donors without previous history of pancreatic disorders were obtained following total pancreatectomy with islet isolation procedure from the University of California, San Francisco and used as controls in this study. Human CP pancreatic tissues of hereditary and idiopathic patients undergoing TPIAT were received from the University of Minnesota. Demographics and characteristics of control and CP subjects, a cohort for single-cell sequencing analysis, were displayed in table 1, online supplemental table 1 and 2. Demographics and characteristics of control and CP subjects, a cohort for the validation experiments of single-cell sequencing data, were displayed in table 2, online supplemental table 4, and 5. All patients with CP provided informed consent, or parental consent and patient assent, as age appropriate. For tissue use in research, the protocol was reviewed and approved by the University of Minnesota and Stanford University Institutional Review Board.

***Patient and public involvement***

Patients or the public were not involved in the design, conduct, reporting, or dissemination plans of this research.

***Tissue processing and single-cell isolation***

To enrich immune cells from human pancreatic tissues, pancreatic tissues were enzymically digested as described in the previous study[1]. After removing red blood cells with RBC lysis buffer (Sigma, R7757), the recovered cells were then washed again with RPMI buffer that contained 10% BCS and suspended in the recovery cell culture freezing medium (Gibco,

#12648-010) to be stored in -80 ºC freezer and liquid nitrogen. To isolate pure leukocytes for the single-cell sequencing analyses, the frozen single cells were thawed and washed twice with RPMI containing 10% FBS. The recovered cells were resuspended in HBSS containing 2% BCS and stained with Live/Dead marker (BioLegend, Zombie Aqua, #423102) and anti-CD45 antibody (BioLegend, PerCP-Cy5.5, #304028) for FACS. FACS was performed with FACSAria IIu at Stanford FACS facility.

***Single-cell CITE-seq and TCR-seq***

Sorted live $CD45^+$ pancreatic immune cells were washed on cold PBS containing 2% BSA and stained with 13 feature antibodies from BioLegend (Totalseq-C, online supplemental table 6) with optimized concentrations. After 30 min incubation with antibodies, cells were washed three times to remove excess antibody. Antibody-labeled cells were loaded onto the Chromium 10x Genomics platform (10x Genomics) to capture single cells with gel beads, which were designed to label single cells for CITE-seq (feature protein expression and 5' whole transcriptomes) and TCR-sequencing according to manufacturer's instructions. Library generation for feature barcoding, gene expression, and V(D)J TCRs was performed using the 10x Chromium Single Cell reagent kits. Following library generation, sequencing was performed on an Illumina HiSeq 4000 (2 x75bp).

***Histology (H&E and Trichome staining) and Immunohistochemistry***

Human pancreatic tissues were fixed with 10% formalin and transferred to 70% ethanol for the paraffin-embedding and further procedures. Paraffin-embedded tissue blocks were sliced and stained for hematoxylin and eosin, Trichrome staining and immunohistochemistry (IHC) with anti-CD45 antibody (Agilent, M070101-2) at Stanford human pathology/histology service center.

***Flow cytometry***

Frozen single cells were thawed and recovered by washing twice with RPMI containing 10% FBS. The recovered cells were resuspended in HBSS containing 2% BCS and stained with flow cytometry antibodies from BioLegend (Live/Dead Zombie Aqua, #423102; anti-CD45, PerCP-Cy5.5, #304028; anti-CD3, Pacific Blue, #300431; anti-CD19, APC-Cy7, #302218; anti-CD4, BV605, #317438; anti-CD8, FITC, #344704; anti-CCR6, APC, #353416). After 30 min incubation with antibodies at RT, cells were washed with HBSS containing 2% BCS and fixed with 4% paraformaldehyde. Flow cytometry was performed with BD LSRII flow cytometer at Stanford FACS facility, and data analysis was performed using Flowjo (BD).

***Chemotaxis assay***

Chemotaxis assay was performed as described in prior studies[2 3]. Immune cells isolated from pancreatic tissues were resuspended at 0.5-1 x $10^6$ cells/ 100 ml in RPMI1640 containing 0.5 % BSA. 600 ml of recombinant human CCL20 (Peprotech, #300-29A) at a final concentration of 500 ng/ml was added to the transwell (Corning, Transwell-24 well with 3mM pore polycarbonate membrane, #3421). 100 ml of cell suspension was added to the upper chamber. Vehicle-added wells were used as negative controls, and CXCL12 (Peprotech, #300-28A) at a final concentration of 100 ng/ml was used as a positive control. The plate was incubated at 37ºC for 3 hours. Total migrating cells were collected from the lower chamber, and total number of cells was assessed by manually counting with hemocytometer. All assays were performed in triplicate.

***Single-cell RNA-seq analysis***

Sample demultiplexing, barcode processing, single-cell counting, and reference genome mapping were performed using the Cell Ranger Software (v3.1.0, GRCh38 ref genome) according to the instruction provided from 10x Genomics. All samples were normalized to present the same effective sequencing depth by using Cell Ranger aggr function. The dimensionality reduction by principal components analysis (PCA), the graph-based clustering and UMAP visualization were performed using Seurat (v3.0, R package). Genes that were detected in less than three cells were filtered out, and cells were filtered out with greater than 10 percent of mitochondrial genes and with fewer than 200 or greater than 50,000 detected genes. We also removed potential doublets by using DoubletFinder (v2.0.2), using 7.5 percentile of the doublet score as cutoff. The differential analysis among clusters were performed using "NormalizeData" and "FindMarkers" functions. The differential test used Wilcoxon ranked-sum method and p value < 0.05 was used for significance after multiple hypothesis correcting by Benjamini & Hochbe method. Cluster annotation was performed based on the top rank of differentially expressed gene list in each cluster and confirmed with the predicted annotations determined by SingleR (v0.99.10, R package).

In order to better characterize the signature of DEGs, we used a fast per-ranked gene set enrichment analysis (GSEA) named fgsea (v1.11.1, R package) to evaluate functional enrichment analysis for DEGs[4]. The hallmark gene sets from Molecular Signatures Database[5] were extracted and used to feed each differentially expressed gene lists. Then adjusted P value < 0.05 were selected as the significantly and functionally enriched biological states or processes.

Receptor–ligand analysis among different groups was performed using CellphoneDB statistical analysis, v.2.0[6]. The significant interactions were selected with p-value < 0.01. Additionally, we also perform the 59 pairs interaction analysis which are common significant pairs identified in all three groups.

#### *Single-cell TCR-seq analysis*

Single-cell TCR clonotypes were assembled using Cell Ranger vdj function. Single-cell barcodes were then used to tie corresponding VDJ (variable-diversity-joining TCR gene segments) and gene expression data simultaneously. Only TCRs with full and paired α and β chain sequences were included in the analysis. TCR Clonotypes were then determined from grouping of cell barcodes that shared the same set of productive CDR3 nucleotide sequences.

Gini coefficient was calculated by using 'repDiversity' function in tcR/immunarch (v0.6.6, R package) with default setting[7].

TCR cluster lineage tracing was performed by considering all clonotypes shared by cells from more than one cluster[8 9]. Raw numbers of cluster clonotype intersections were analyzed and visualized as upset plots.

#### *GLIPH2 analysis*

Paired CDR3β and CDR3α sequences from pancreatic T cells across control, hereditary and idiopathic CP were collected and analyzed using GLIPH2 downloaded from http://50.255.35.37:8080/. Default parameters and human v2.0 reference dataset were used for this analysis. Candidates clusters were selected from the motifs with final score less than $10^{-8}$ and shared by at least two different individuals.

***Single-cell sequencing with feature antibody staining (CITE-seq)***

Feature barcode (10X genomics) was used to link the analysis of gene expression and cell surface protein simultaneously in the same cell. 10 CITE–seq samples were demultiplexed, barcode processing, generating protein expression profile using Cell Ranger Software. A filtered feature counts matrix was then imported in the Seurat to extract protein panel by using 'Antibody Capture' function and to do following exploratory analysis. To better compare the protein data with transcriptomic data, we first use the cluster identified in the above described 'scRNA-seq analysis' to draw featureplot and heatmap. Then we reclustered based on CITE–seq protein panel. Protein expression was first normalized and euclidean distances between cells were then computed from normalized protein expression of all features (10 antibody readouts) using the dist function in R, with default parameters. UMAP clustering was run using distance matrix defined as input with resolution of 0.5.

***Statistical analysis***

Unless otherwise stated, statistical comparisons were performed by using GraphPad software (Prism, version 8.1.0). Two groups comparisons were performed with two-tailed Student's t-test, and three groups comparisons were done with one-way ANOVA with Tukey's multiple comparison test or Kruskal-Wallis test. Differences were considered to be statistically significant when $p<0.05$.

**Supplemental Figures.**

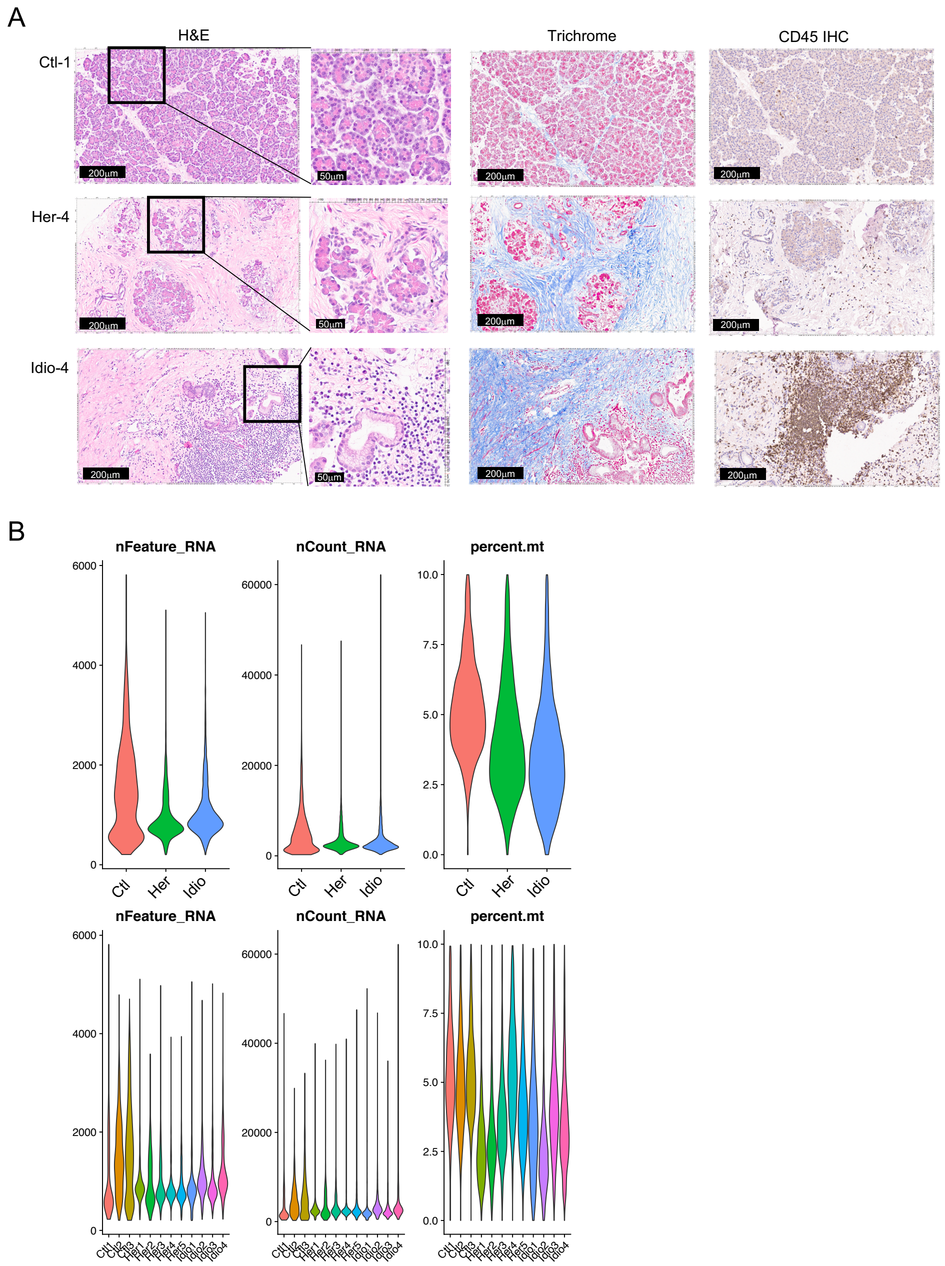


**Supplemental Figure 1. Histology of pancreas tissues from controls and CP.** **(A)** H&E, Trichrome, and immunohistochemistry staining of pan-leukocyte marker, CD45 with representative pancreas tissues from control (#1), hereditary CP (#4), and idiopathic CP (#4). **(B)** Quality control of single-cell sequencing data. Cells were filtered out with greater than 10% of mitochondrial genes and with fewer than 200 or with greater than 50,000 detected genes (nFeature, number of transcripts; nCount, number of reads; percent.mt, % of mitochondrial genes).

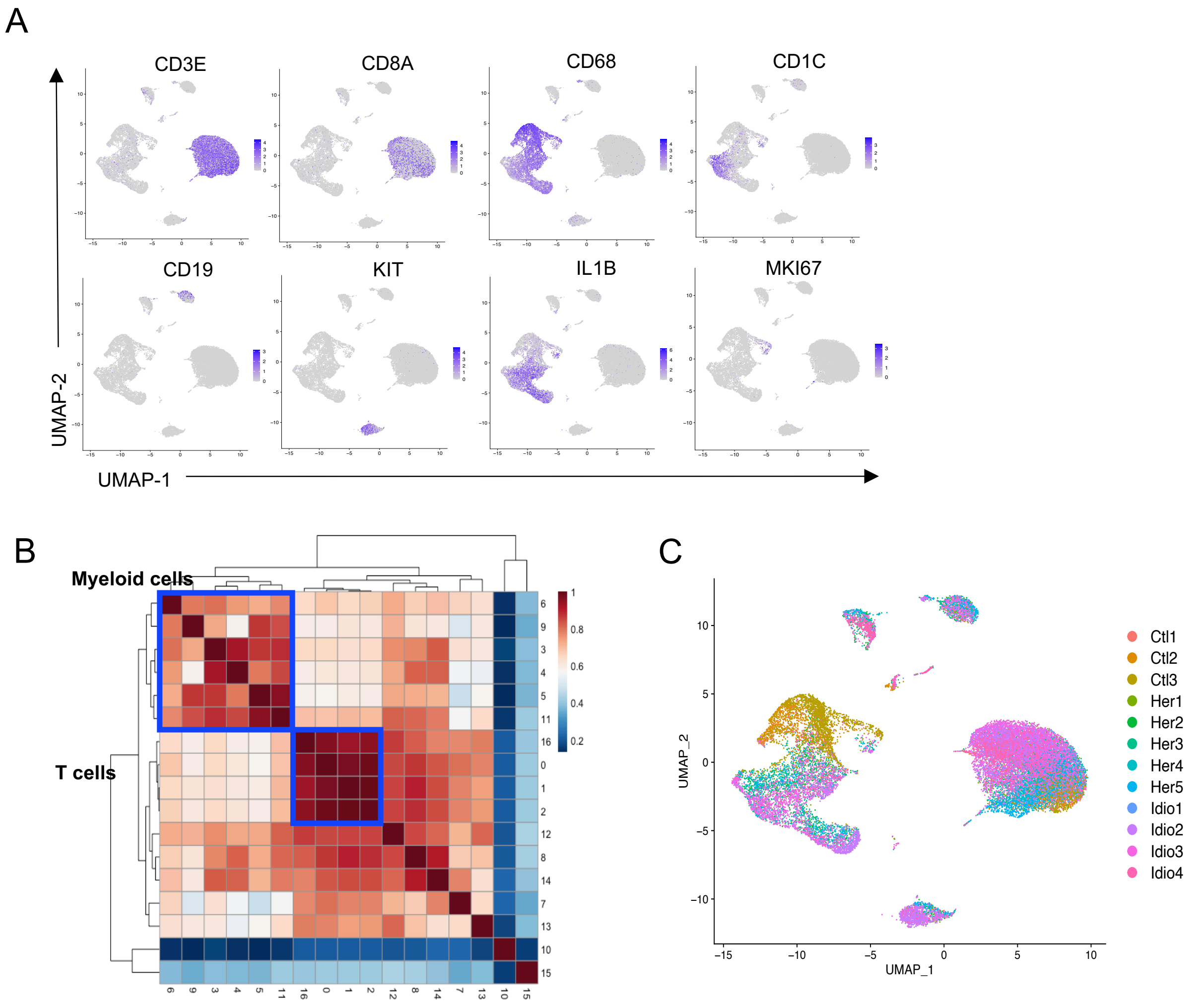


**Supplemental Figure 2. Single-cell transcriptome analysis of human pancreatic immune cells from organ donors and CP patients.** (**A**) UMAP plots of immune cell marker gene expressions in total immune cells; *CD3E, CD8A* for T cells; *CD68* for macrophages; *CD1C* for dendritic cells; *CD19* for B cells; *KIT* for mast cells; *IL1B* for pro-inflammatory myeloid cells; *MKI67* for proliferating cells. (**B**) Heatmap of correlation matrix among 17 pancreatic immune clusters identified by clustering analysis. (**C**) UMAP plot of total pancreatic immune cells colored by individual subjects.

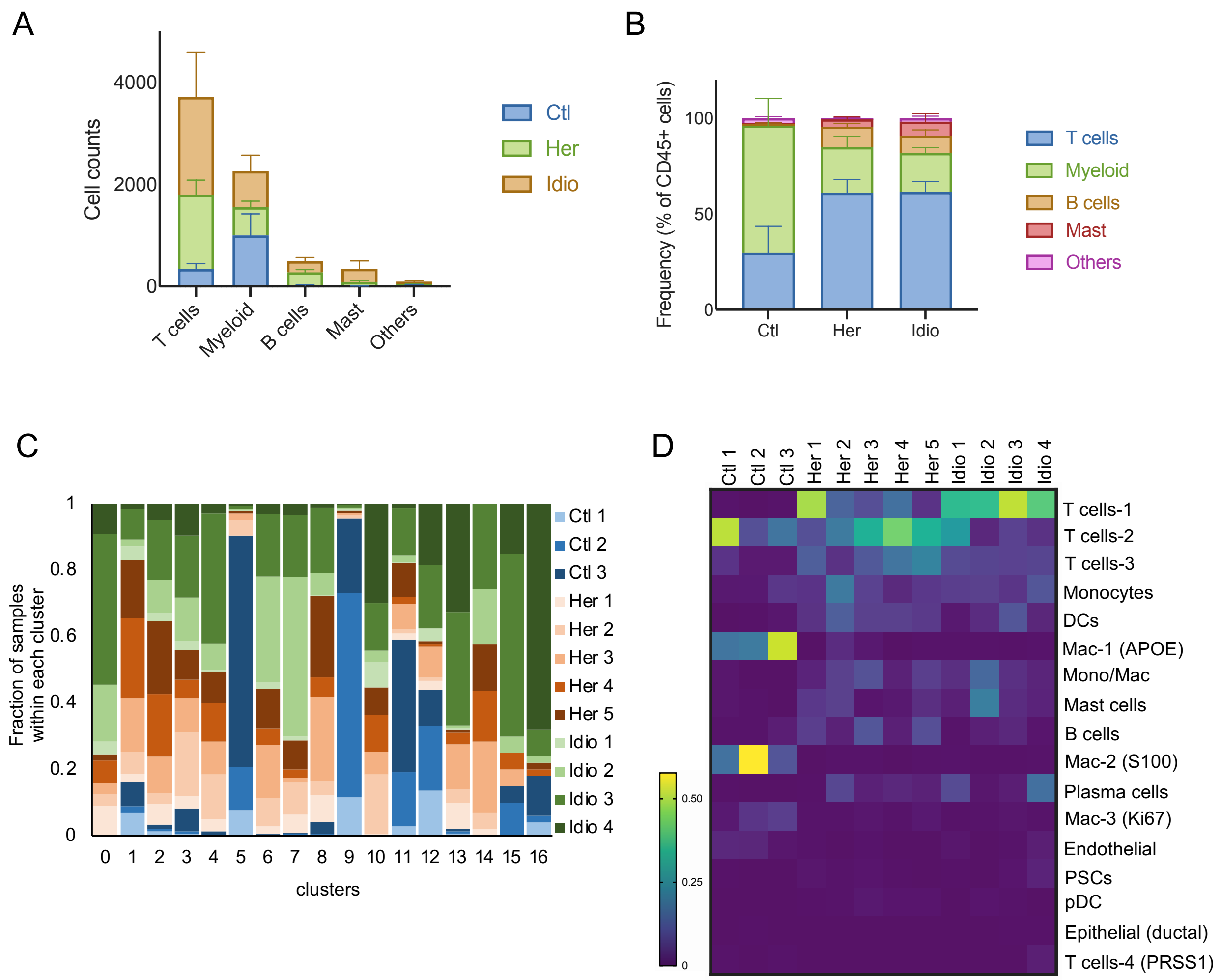


**Supplemental Figure 3. Composition of human pancreatic immune populations from organ donors and CP patients.** (**A**) Bar graph displaying major pancreatic immune populations by cell counts colored based on the group. (**B**) Bar graph showing the composition of immune cells in each group colored based on immune subpopulations. (**C**) Subjects' contribution to the cluster shown as fractions of 12 subjects in each cluster identified by clustering analysis. (**D**) Heatmap of immune cluster frequencies in controls and CP subjects.

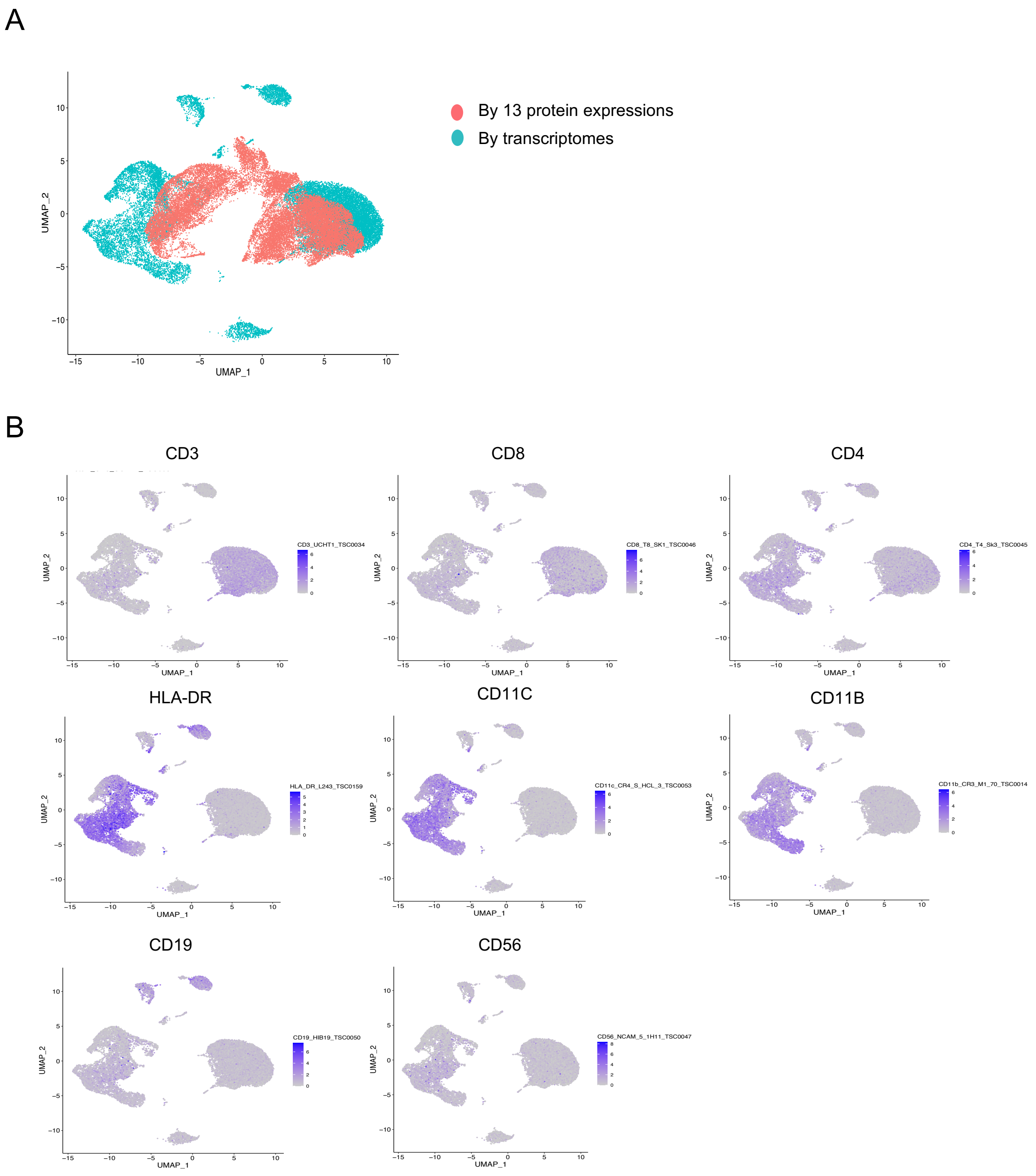


**Supplemental Figure 4. CITE-seq analysis of human pancreatic immune cells from organ donors and CP patients.** (**A**) UMAP plots of protein expressions (across 10 samples; 3 controls, 5 hereditary CP, and 2 idiopathic CP) and transcriptomes (across 12 samples; 3 controls, 5 hereditary CP, and 4 idiopathic CP). (**B**) UMAP plots of representative immune marker protein expressions; CD3, CD8, and CD4 for T cells; HLA-DR, CD11C, and CD11B for myeloid cells; CD19 for B cells; CD56 for NK cells.

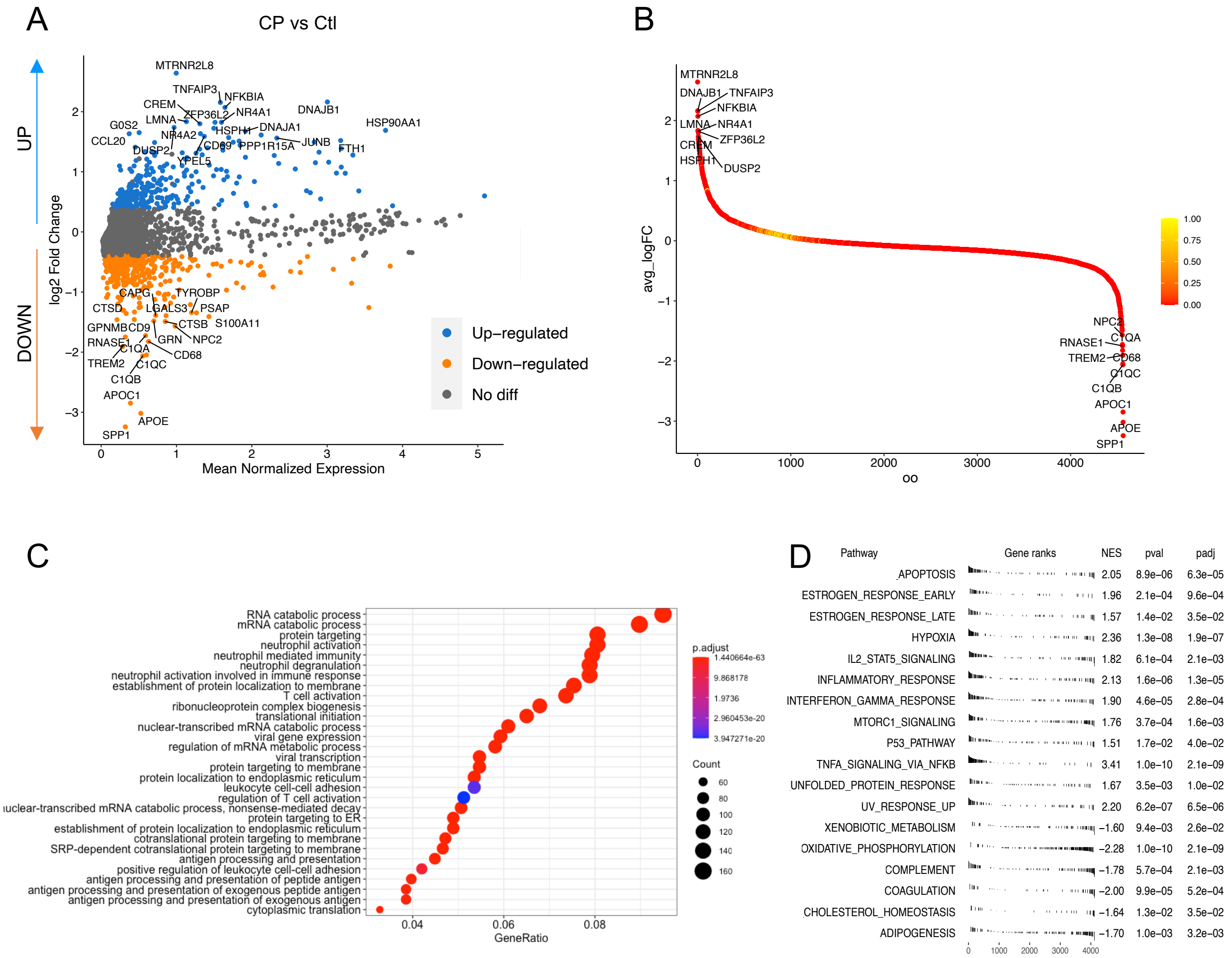


**Supplemental Figure 5. DEGs and functional enrichment analysis of pancreatic immune cells from controls and CP.** (**A**) Volcano plot of DEG analysis between control and CP with total immune populations. Each dot represents a gene, with significantly up-regulated top 20 genes and down-regulated top 20 genes in CP versus control colored blue and yellow, respectively. (**B**) DEGs ranked by fold changes in CP versus control. (**C**) Functional enrichment analysis displaying biological processes enriched in CP versus control. (**D**) Functional enrichment analysis showed significantly enriched pathways in CP versus control.

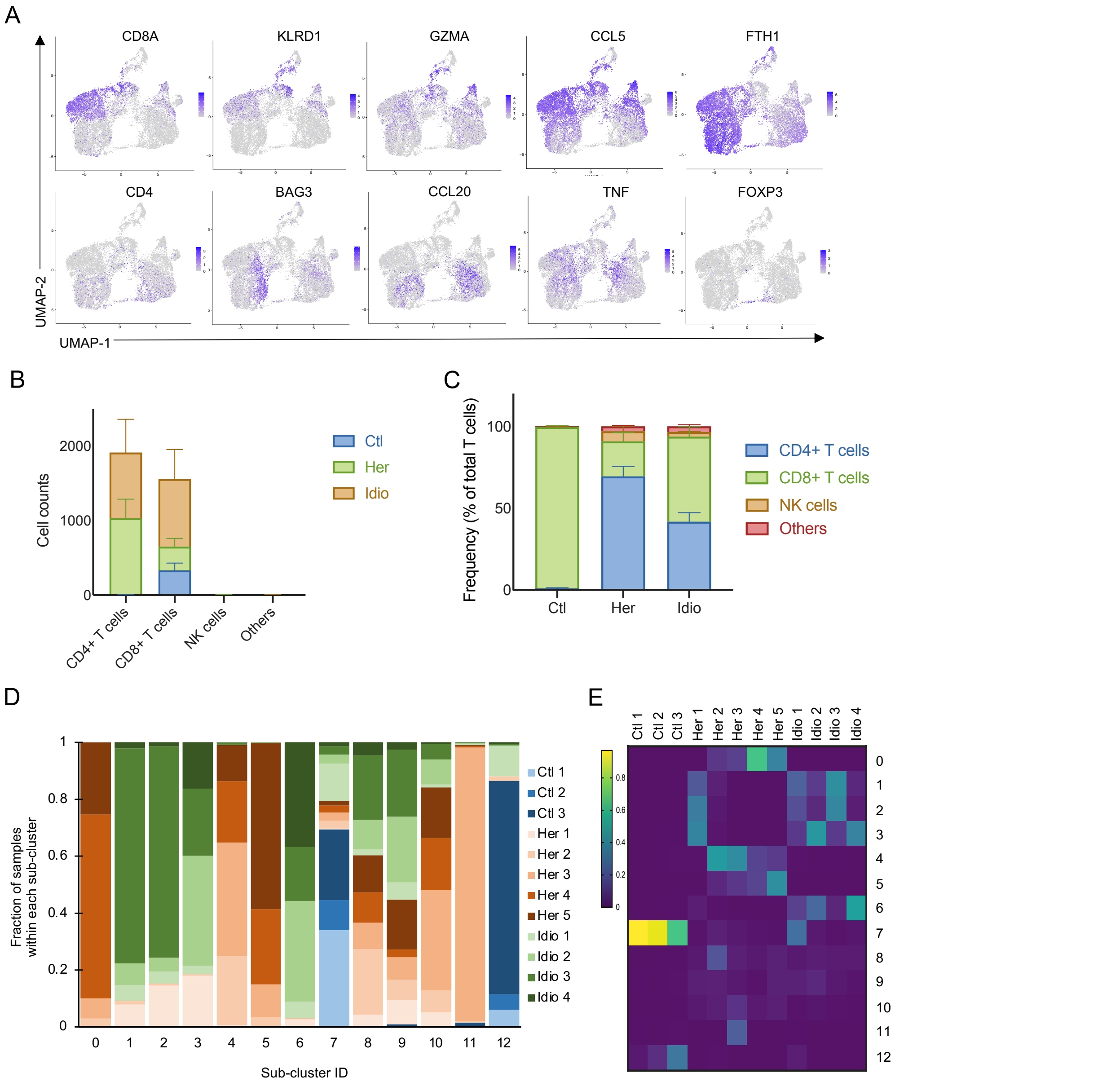


**Supplemental Figure 6. Single-cell transcriptome analysis of human pancreatic T cells from organ donors and CP patients.** (**A**) UMAP plots of significantly expressed genes in T cell subclusters and their expression patterns. (**B**) Bar graph displaying major pancreatic T cell subpopulations by cell counts colored based on the group. (**C**) Bar graph showing the composition of T cell subpopulations in each group colored based on immune subpopulations. (**D**) Subjects' contribution to the cluster shown as fractions of 12 subjects in each cluster identified by clustering analysis. (**E**) Heatmap of T cell subcluster frequencies in controls and CP subjects.

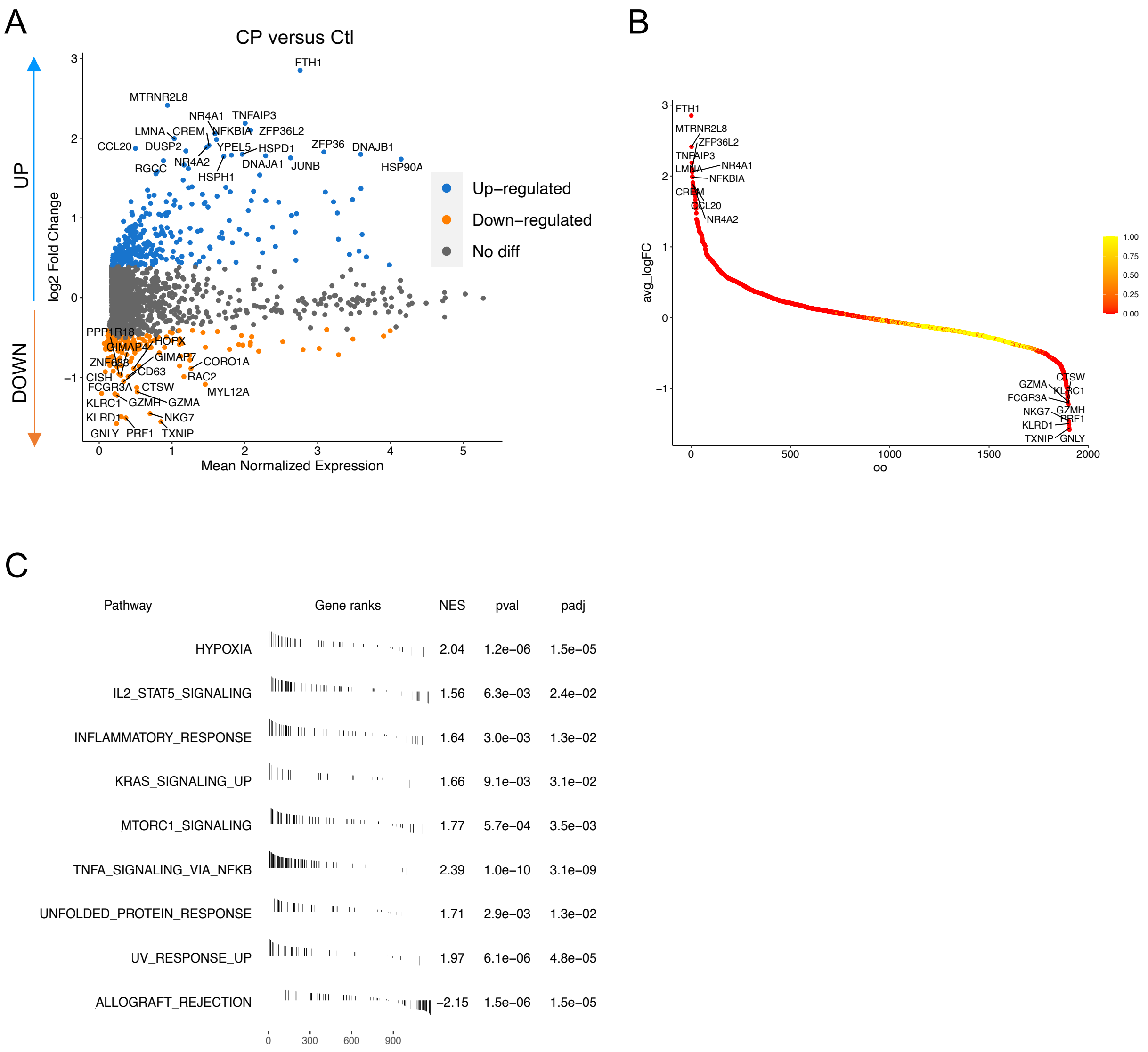


**Supplemental Figure 7. DEGs and functional enrichment analysis of pancreatic T cells from controls and CP.** (**A**) Volcano plot of DEG analysis between control and CP with T cell populations. Each dot represents a gene, with significantly up-regulated top 20 genes and down-regulated top 20 genes in CP versus control colored blue and yellow, respectively. (**B**) DEGs ranked by fold changes in CP versus control. (**C**) Functional enrichment analysis showed significantly enriched pathways in CP versus control.

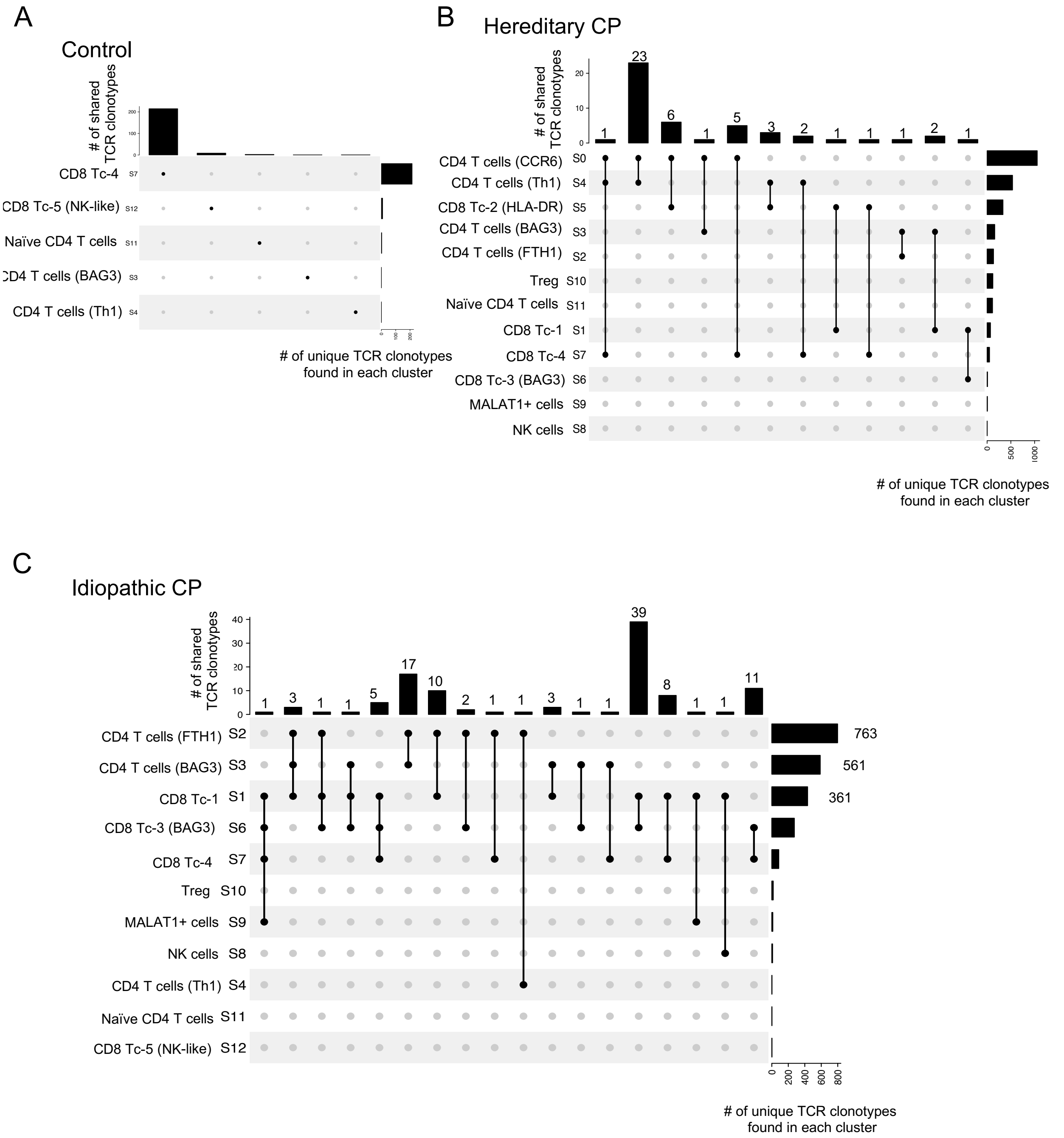


**Supplemental Figure 8. Single-cell transcriptome and TCR analyses of human pancreatic T cells trace T cell lineages and interactions among different T cell subclusters.** Upset plots displaying TCR clonotypes shared among T cell clusters in control (**A**), hereditary CP (**B**), and idiopathic CP (**C**). Each shared clonotype was indicated by black dots with a connected black line. The horizontal bar graph indicates the total number of shared TCR clonotypes for cluster intersections, and the vertical bar graph indicates the clonotype count in a single cluster.

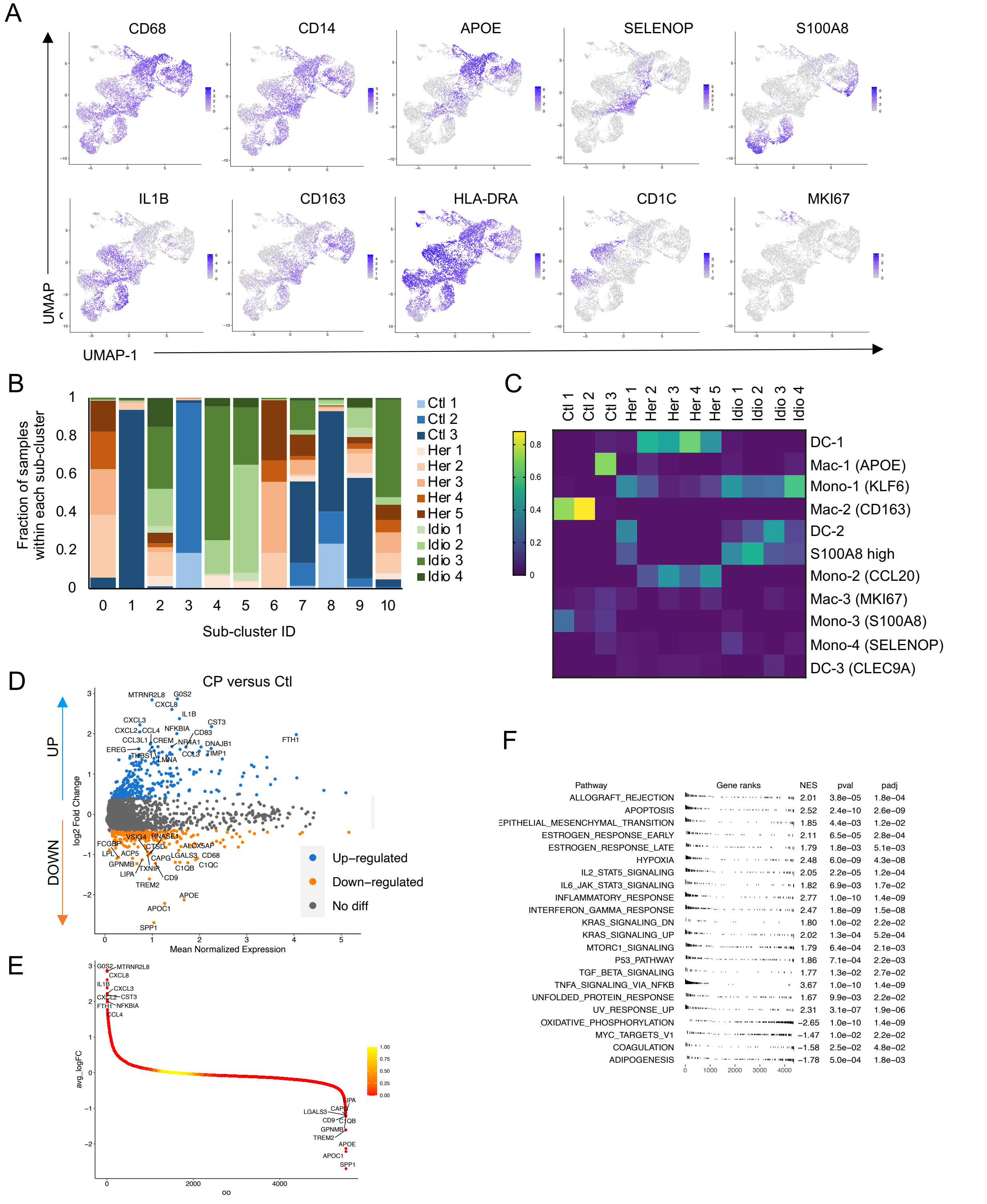


**Supplemental Figure 9. Single-cell transcriptome analysis of human pancreatic myeloid cells from organ donors and CP patients.** (**A**) UMAP plots of significantly expressed genes in myeloid cell subclusters and their expression patterns. (**B**) Subjects' contribution to the cluster shown as fractions of 12 subjects in each cluster identified by clustering analysis. (**C**) Heatmap of myeloid cell subcluster frequencies in controls and CP subjects. (**D**) Volcano plot of DEG analysis between control and CP with myeloid cell populations. Each dot represents a gene, with significantly up-regulated top 20 genes and down-regulated top 20 genes in CP versus control colored blue and yellow, respectively. (**E**) DEGs ranked by fold changes in CP versus control. (**F**) Functional enrichment analysis showed significantly enriched pathways in CP versus control.

**Supplemental tables.**

| | Group | Genetic mutations | Age | Gender | BMI | Severity score | Disease duration (years) |
|---|---|---|---|---|---|---|---|
| 1 | Control | - | 55 | M | 22.9 | NA | NA |
| 2 | | - | 35 | F | 33.3 | NA | NA |
| 3 | | - | 43 | F | 22 | NA | NA |
| 1 | Her | PRSS1 | 21 | M | 23.1 | 9 | 17 |
| 2 | | PRSS1 | 9 | F | 20.3 | 8 | 7 |
| 3 | | PRSS1, CFTR, SPINK1 | 15 | M | 21.3 | 10 | 2 |
| 4 | | PRSS1 | 13 | M | 18.2 | 9 | 10 |
| 5 | | PRSS1, CFTR | 16 | F | 22 | 9 | 7 |
| 1 | Idio | none | 47 | F | 19.1 | 1 | 4 |
| 2 | | none | 42 | F | 27.3 | 9 | 3 |
| 3 | | none | 31 | F | 28.8 | 6 | 3 |
| 4 | | none | 42 | F | 28.7 | 5 | 17 |

**Supplemental Table 1. Demographic and characteristics of control and CP subjects, a cohort for single-cell sequencing analysis.**

| Characteristics | Hereditary (n=5) | Idiopathic (n=4) | P value |
|---|---|---|---|
| Age (years, mean ± SD) [A] | 14.8 ± 4.4 | 40.5 ± 6.8 | **0.0002** |
| Sex, m/f [B] | 3/2 | 0/4 | 0.06 |
| Height (meters, mean ± SD) [A] | 1.62 ± 0.18 | 1.62 ± 5.4 | 0.98 |
| Weight (kg, mean ± SD) [A] | 56.1 ± 15.5 | 68.4 ±15.6 | 0.28 |
| Body mass index (kg/$m^2$, mean ± SD) [A] | 21.0 ± 1.9 | 26 ± 4.6 | 0.06 |
| Severity gross fibrosis score [A] (median, IQR) | 9 [8-10] | 5.5 [1-9] | **0.04** |
| Pancreatitis Duration [A] (years, mean ± SD) | 8.6 ± 5.6 | 6.9 ± 7.2 | 0.71 |
| HbA1c (%, mean ±SD) [A] | 5.4 ±0.3 | 5.1 ±0.6 | 0.37 |
| Glucose (mg/dL, mean ±SD) [A] | 94.0 ±9.4 | 89.5 ±14.5 | 0.59 |
| MRCP use [B] | 5 (100%) | 4 (100%) | - |
| Atrophy by MRI [B] | 3 (60%) | 0 (0%) | 0.06 |
| Stone or Calcification by MRI [B] | 1 (20%) | 1 (25%) | 0.86 |
| Dilated Pancreatic Duct by MRI [B] | 5 (100%) | 1 (25%) | **0.02** |
| Side Branches by MRI [B] | 3 (60%) | 1 (25%) | 0.29 |

SD, standard deviation; IQR, interquartile range.

[A]Unpaired two-tailed t-test; [B]Chi-square was used for p-values, Bold p value, $p<0.05$.

**Supplemental Table 2. Statistical comparisons of demographic and characteristics between hereditary and idiopathic CP patients, a cohort for single-cell sequencing analysis.**

| | Group | Genetic mutations | Transcriptome | Surface Antibody | TCR | Cell count (Before QC) | Cell count (After QC) | Doublet count | Single cells | Group cell count | Total cell count |
|---|---|---|---|---|---|---|---|---|---|---|---|
| 1 | Control | - | O | O | O | 939 | 808 | 61 | 747 | 4169 | 28547 |
| 2 | | - | O | O | O | 1525 | 1246 | 93 | 1153 | | |
| 3 | | - | O | O | O | 3225 | 2453 | 184 | 2269 | | |
| 1 | Her | PRSS1 | O | O | O | 1539 | 1471 | 110 | 1361 | 11786 | |
| 2 | | PRSS1 | O | O | O | 2367 | 2126 | 159 | 1967 | | |
| 3 | | PRSS1, CFTR, SPINK1 | O | O | O | 3010 | 2814 | 211 | 2603 | | |
| 4 | | PRSS1 | O | O | O | 3806 | 3250 | 244 | 3006 | | |
| 5 | | PRSS1, CFTR | O | O | O | 3437 | 3080 | 231 | 2849 | | |
| 1 | Idio | none | O | O | O | 941 | 844 | 63 | 781 | 12592 | |
| 2 | | none | O | X | O | 3826 | 3752 | 281 | 3471 | | |
| 3 | | none | O | O | O | 7670 | 7223 | 542 | 6681 | | |
| 4 | | none | O | X | O | 2270 | 1793 | 134 | 1659 | | |

**Supplemental Table 3. Single-cell sequencing data information (O, applied; X, not applied).**

| | Group | Genetic mutations | Age | Gender | BMI | Severity score | Disease duration (years) |
|---|---|---|---|---|---|---|---|
| 1 | Ctl | - | 38 | F | 32.1 | NA | NA |
| 2 | | - | 25 | M | 24.8 | NA | NA |
| 3 | | - | 33 | M | 34.4 | NA | NA |
| 4 | | - | 48 | F | 30.7 | NA | NA |
| 5 | | - | 26 | F | 25.3 | NA | NA |
| 6 | | - | 29 | M | 28.9 | NA | NA |
| 7 | | - | 19 | M | 30.7 | NA | NA |
| 1 | Her | CFTR, CTRC | 41 | F | 18.1 | 9 | 2.67 |
| 2 | | SPINK1 | 20 | F | 31.1 | 9 | 4.33 |
| 3 | | PRSS1, CFTR, SPINK1 | 15 | M | 21.3 | 10 | 2.00 |
| 4 | | CFTR, SPINK1 | 19 | M | 24.4 | 5 | 3.83 |
| 5 | | SPINK1 | 14 | M | 19.1 | 9 | 12.17 |
| 6 | | CFTR | 20 | F | 21.3 | 8 | 2.17 |
| 7 | | PRSS1, SPINK1 | 20 | M | 30.9 | 9 | 9.00 |
| 8 | | CFTR | 29 | M | 23.5 | 9 | 9.17 |
| 1 | Idio | none | 50 | F | 28.3 | 2 | 2.75 |
| 2 | | none | 47 | F | 19.1 | 1 | 4.50 |
| 3 | | none | 31 | M | 34 | 6 | 24.42 |
| 4 | | none | 24 | F | 27 | 5 | 3.42 |

**Supplemental Table 4. Demographic and characteristics of control and CP subjects, a cohort for functional validation analyses.**

| Characteristics | Hereditary (n=8) | Idiopathic (n=4) | P value |
|---|---|---|---|
| Age (years, mean ± SD) [A] | 22.3 ± 8.8 | 38.0 ± 12.5 | **0.029** |
| Sex, m/f [B] | 5/3 | 1/3 | 0.22 |
| Height (meters, mean ± SD) [A] | 1.70 ± 6.8 | 1.62 ± 11.9 | 0.21 |
| Weight (kg, mean ± SD) [A] | 68.6 ± 16.9 | 73.5 ±27.0 | 0.71 |
| Body mass index (kg/m$^2$, mean ± SD) [A] | 23.7 ± 4.9 | 27.1 ± 6.1 | 0.32 |
| Severity gross fibrosis score [A] (median, IQR) | 9 [8-10] | 5.5 [1-9] | **0.001** |
| Pancreatitis Duration [A] (years, mean ± SD) | 5.7 ± 3.9 | 8.8 ± 10.5 | 0.46 |
| HbA1c (%, mean ±SD) [A] | 5.3 ±0.3 | 5.3 ±0.3 | 0.90 |
| Glucose (mg/dL, mean ±SD) [A] | 91.4 ± 4.3 | 84.5 ±7.8 | 0.07 |
| MRCP use [B] | 8 (100%) | 3 (75%) | - |
| Atrophy by MRI [B] | 3 (37.5%) | 1 (25%) | 0.90 |
| Stone or Calcification by MRI [B] | 2 (25%) | 0 (0%) | 0.34 |
| Dilated Pancreatic Duct by MRI [B] | 6 (75%) | 1 (25%) | 0.20 |
| Side Branches by MRI [B] | 3 (37.5%) | 0 (0%) | 0.21 |

SD, standard deviation; IQR, interquartile range.

[A]Unpaired two-tailed t-test; [B]Chi-square was used for p-values, Bold p value, $p<0.05$.

**Supplemental Table 5. Statistical comparisons of demographic and characteristics between hereditary and idiopathic CP patients, a cohort for functional validation analyses.**

| | **Antibody information (BioLegends)** |
|---|---|
| 1 | 368545 TotalSeq™-C0048 anti-human CD45 (clone: 2D1) |
| 2 | 300479 TotalSeq™-C0034 anti-human CD3 |
| 3 | 302265 TotalSeq™-C0050 anti-human CD19 |
| 4 | 344651 TotalSeq™-C0045 anti-human CD4 |
| 5 | 344753 TotalSeq™-C0046 anti-human CD8 |
| 6 | 362559 TotalSeq™-C0047 anti-human CD56 (NCAM) |
| 7 | 302649 TotalSeq™-C0085 anti-human CD25 |
| 8 | 371521 TotalSeq™-C0053 anti-human CD11c |
| 9 | 101275 TotalSeq™-C0014 anti-mouse/human CD11b |
| 10 | 321147 TotalSeq™-C0205 anti-human CD206 (MMR) |
| 11 | 307663 TotalSeq™-C0159 anti-human HLA-DR |
| 12 | 353251 TotalSeq™-C0148 anti-human CD197 (CCR7) |
| 13 | 304163 TotalSeq™-C0063 anti-human CD45RA |

**Supplemental Table 6. Feature antibodies used for CITE-seq analysis.**

### References

1. Lee B, Adamska JZ, Namkoong H, et al. Distinct immune characteristics distinguish hereditary and idiopathic chronic pancreatitis. *J Clin Invest* 2020;130(5):2705-11. doi: 10.1172/JCI134066 [published Online First: 2020/02/14]

2. Facco M, Baesso I, Miorin M, et al. Expression and role of CCR6/CCL20 chemokine axis in pulmonary sarcoidosis. *J Leukoc Biol* 2007;82(4):946-55. doi: 10.1189/jlb.0307133 [published Online First: 2007/07/07]

3. Francis JN, Sabroe I, Lloyd CM, et al. Elevated CCR6+ CD4+ T lymphocytes in tissue compared with blood and induction of CCL20 during the asthmatic late response. *Clin Exp Immunol* 2008;152(3):440-7. doi: 10.1111/j.1365-2249.2008.03657.x [published Online First: 2008/04/22]

4. Korotkevich G, Sukhov V, Sergushichev A. Fast gene set enrichment analysis. *bioRxiv* 2019

5. Liberzon A, Birger C, Thorvaldsdottir H, et al. The Molecular Signatures Database (MSigDB) hallmark gene set collection. *Cell Syst* 2015;1(6):417-25. doi: 10.1016/j.cels.2015.12.004 [published Online First: 2016/01/16]

6. Efremova M, Vento-Tormo M, Teichmann SA, et al. CellPhoneDB: inferring cell-cell communication from combined expression of multi-subunit ligand-receptor complexes. *Nat Protoc* 2020;15(4):1484-506. doi: 10.1038/s41596-020-0292-x [published Online First: 2020/02/28]

7. Nazarov VI, Pogorelyy MV, Komech EA, et al. tcR: an R package for T cell receptor repertoire advanced data analysis. *BMC Bioinformatics* 2015;16:175. doi: 10.1186/s12859-015-0613-1 [published Online First: 2015/05/29]

8. Lex A, Gehlenborg N, Strobelt H, et al. UpSet: Visualization of Intersecting Sets. *IEEE Trans Vis Comput Graph* 2014;20(12):1983-92. doi: 10.1109/TVCG.2014.2346248 [published Online First: 2015/09/12]

9. Corridoni D, Antanaviciute A, Gupta T, et al. Single-cell atlas of colonic CD8(+) T cells in ulcerative colitis. *Nat Med* 2020;26(9):1480-90. doi: 10.1038/s41591-020-1003-4 [published Online First: 2020/08/05]
